# Anticodon Edited Transfer RNAs (ACE-tRNAs) Encoded as Therapeutic Nonviral Minimal DNA Vectors

**DOI:** 10.1101/2025.09.06.674645

**Authors:** Joseph J. Porter, Wooree Ko, Emily G. Sorensen, Zachary Cheung, Tyler Couch, Jeffrey T. Gabell, Victoria Shwe, Julia Hyatt, Jennasea B. Licata, Luke K. Peterson, David A. Dean, John D. Lueck

**Author notes:** These authors contributed equally (Joseph J. Porter, Wooree Ko).

## Abstract

Nonsense mutations, resulting from a premature termination codon (PTC), make up ∼11% of all genetic lesions causing disease, affecting millions of people worldwide. Nonsense suppressor anticodon-edited tRNAs (ACE-tRNAs) have emerged as a therapeutic modality for the rescue of PTCs. Delivery of ACE-tRNAs *in vivo* has been achieved by adeno-associated viral vector and RNA-lipid nanoparticle, however due to drawbacks associated with these approaches, DNA delivery remains an attractive approach. DNA-based approaches afford ease of manufacturing at a relatively low cost and exhibit improved therapeutic durability and safety as compared to viral vector-or RNA-based approaches. Due to the small size of human tRNA genes employed as ACE-tRNAs, in principle, DNA vectors <200 base pairs (bp) in size (minivectors) could be utilized for delivery of actively transcribed ACE-tRNAs. Here we demonstrate that linear DNA ACE-tRNA vectors as small as 200 bp effectively suppress several nonsense mutations in *CFTR* and *REP1*, and that ACE-tRNA minivectors display significantly improved bioavailability, reduced innate immune burden, and superior biostability as compared to conventional plasmid DNA vectors.

## INTRODUCTION

Nonsense mutations result from single nucleotide DNA variants that convert an amino-acid-encoding codon to one of the three stop codons (UAA, UAG, UGA). Due to the presence of a stop codon within the protein coding region of mRNA transcripts, these mutations are also referred to as premature termination codons (PTCs). Introduction of a PTC results in little-to-no full-length protein production from the affected gene, often with deleterious cellular effects. To that end, mutations resulting in PTCs lead to more than 700 human diseases (1), accounting for ∼10% of all incidences of genetic disease (2), with development of viable therapeutic approaches giving the potential for treatment to millions of people worldwide. Loss-of-protein results both from premature translation termination leading to production of truncated protein products, and from loss of PTC-containing mRNA transcript due to the action of the nonsense mediated decay (NMD) quality control pathway (3).

Although PTCs present a unifying mechanism of genetic disease upon which considerable efforts have been focused, to date no specific therapy for PTCs has reached widespread clinical use. Therapeutic approaches under development for rescue of PTCs include CRISPR/Cas approaches (4), mRNA pseudouridylation (5), antisense oligo (ASO) (6), small molecule translational readthrough inducing drugs (TRIDs) (7), and nonsense suppressor tRNAs (sup-tRNAs) also known as anticodon-edited tRNAs (ACE-tRNAs) (8–10). The ACE-tRNA approach has long been envisioned for the clinical rescue of PTCs (11), with development of this approach enjoying a resurgence in enthusiasm following the demonstration of safety and efficacy in cell and mouse models of PTC-associated diseases (12–14). While the primary safety concern for therapeutic application of ACE-tRNAs has been readthrough of natural termination codons (NTCs), all investigations to date have demonstrated significant suppression of PTCs with minimal NTC readthrough (12–14).

The ACE-tRNA as a therapeutic modality has been delivered using several different approaches (Fig. S1A) including direct RNA delivery (12,14,15), expression from a DNA vector (12,15–21), and expression from viral vectors (13,16,22). Direct delivery of the ACE-tRNA as RNA in lipid nanoparticle formulations has demonstrated safety and efficacy in a mouse model of cystic fibrosis (CF) (14), however the RNA has been demonstrated to have a relatively short half-life (15), with frequent redosing potentially causing cytotoxicity due to accumulated application of the nanoparticle carrier lipid or polymer (23). Adeno-associated viral vectors (AAVs) have achieved clinical success with FDA approved total gene replacement therapies for lebers congenital amaurosis (24), spinal muscular atrophy (25), and Duchenne muscular dystrophy (26). Viral vectors have evolved effective cell entry, endosomal escape, intracellular trafficking, and nuclear entry capabilities unmatched by current human-engineered nucleotide delivery efforts (27,28). However, interest in AAVs as a therapeutic vector is somewhat tempered by issues related to cost of manufacturing, the ∼5 kb upper limit of transgene packaging capacity, uncertain duration of effect, inability for redosing, and the existence of patient populations with pre-existing immunity to therapeutic AAVs (27,29–32). Expression of ACE-tRNAs from plasmid-based DNA vectors has demonstrated efficacy in both human cell culture and mouse models (12,15–21).

Therapeutic plasmid DNA (pDNA) vectors are composed of a region containing a therapeutic cargo, generally a gene expression cassette, and the plasmid backbone containing a bacterial origin of replication and antibiotic resistance marker for selection (Fig. S1B). While the DNA backbone components are indispensable for pDNA replication in bacteria, the pDNA backbone elements lead to silencing of the therapeutic cargo, and in the case of ACE-tRNAs, occupy a dramatically larger portion of the DNA vector than the therapeutic cargo (Fig. S1B).

Due to drawbacks associated with the use of DNA plasmids as therapeutic vectors, several DNA “minivector” production approaches have been developed (33–35).

Here we demonstrate that ACE-tRNA minivectors of varying DNA size and shape can be produced and are competent to rescue clinically relevant levels of several PTC-containing genes for proteins associated with human diseases including cystic fibrosis (CF) and choroideremia (CHM). We demonstrate that DNA vectors as small as 200 bp support effective ACE-tRNA transcription (Fig. S1C) resulting in PTC rescue, while also displaying better bioavailability and stability than pDNA therapeutic vectors. Functional rescue of PTCs with ACE-tRNA minivectors opens up new potential avenues for the development of therapeutic approaches for the rescue of PTC-containing protein expression associated with many genetic diseases.

## METHODS

Unless otherwise noted, all oligonucleotides were obtained from Integrated DNA Technologies (IDT) and all molecular biology reagents were obtained from New England Biolabs (NEB). All pDNA was prepared by Nucleobond Xtra Midi EF kit (Macherey-Nagel).

### Expression and purification of Pfu-Sso7d polymerase

NiCo(DE3) cells (NEB) containing a plasmid expressing Pfu-Sso7d polymerase (121) were grown overnight in ZYM-505 media (122), induced with 1 mM IPTG, grown at 37 °C for 4 hours, and the resulting protein was purified from the *E. coli* lysate by Ni-NTA IMAC. The purified polymerase was buffer exchanged using PD-10 desalting columns (GE Healthcare) into storage buffer (100 mM Tris-HCl, pH 8.0, 0.2 mM EDTA, 0.2% NP-40 substitute, 0.2% Tween 20, freshly dissolved 2 mM DTT, and 50% glycerol final concentration). The resulting enzymatic polymerase activity was titrated using a 2-fold serial dilution and the dilution giving the highest PCR product yield was used to dilute the rest of the stock, which was stored at-80 °C.

### Expression and purification of T4 DNA ligase

A DNA fragment containing the T4 DNA ligase sequence from Addgene plasmid #87742 (123) was cloned into a pET21a vector (see supplemental information for plasmid DNA sequence) by Gibson assembly (NEBuilder HiFi mix). Protein from the resulting expression vector was expressed and purified as outlined above for the Pfu-Sso7d polymerase. The purified T4 DNA ligase was buffer exchanged into storage buffer (10 mM Tris-HCl, pH 7.4, 50 mM KCl, 1 mM freshly dissolved DTT, 0.1 mM EDTA, and 50% glycerol), diluted to ∼400 units/μL (as compared to NEB T4 DNA ligase, M0202S), aliquoted, and stored at-80 °C.

### Expression and purification of TFAM

The DNA binding domain from transcription factor A, mitochondrial (TFAM) was synthesized (Genscript) and expressed from a pET28a(+) vector (see supplemental information for plasmid DNA sequence) and purified as outlined above (Fig. S2). The purified TFAM was exchanged into storage buffer (25 mM Tris-HCl, pH 8, 100 mM NaCl, freshly dissolved 1 mM DTT, and 5% glycerol), aliquoted, and stored at-80 °C. The TFAM protein concentration was determined by A_280_ using a molar extinction coefficient of 42,450 M^-1^ * cm^-1^ and a molecular mass of 31.4 kDa.

### Expression and purification of TelN

The protelomerase from phage N15 (TelN) was amplified from the pJAZZ-OK plasmid (Lucigen) and was cloned into a pET21a vector (see supplemental information for plasmid DNA sequence) by Gibson assembly (NEBuilder HiFi mix). TelN was expressed and purified as outlined above (Fig. S4). An additional cation exchange purification step was added to the TelN purification scheme. TelN was eluted from the Ni-NTA column in low salt elution buffer (50 mM Tris-HCl, pH 8, 10 mM NaCl, 500 mM imidazole, 10% glycerol), bound to a HiTrap SP column, washed with 10 resin bed volumes of ion exchange wash buffer (50 mM Na_2_HPO_4_, pH 7.6, 10 mM NaCl, 0.5 mM EDTA, 1 mM DTT, and 10% glycerol), and eluted with ion exchange elution buffer (50 mM Na_2_HPO_4_, pH 7.6, 500 mM NaCl, 0.5 mM EDTA, 1 mM DTT, and 10% glycerol). The purified TelN was buffer exchanged into storage buffer (20 mM Tris-HCl, pH 7.5, 250 mM potassium acetate, 0.1 mM EDTA, 1 mM DTT, 2% Triton X-100, 50% glycerol), aliquoted, and stored at-80 °C. The TelN activity was titrated using a plasmid containing 2 *telRL* recognition sites, with cleavage products assayed by agarose gel electrophoresis. A typical concentration of TelN activity was ∼20 units/μL, with one unit defined as the amount of enzyme required to cleave 313 fmol of *telRL* sites, in a total reaction volume of 50 μL in 30 minutes at 30 °C in 1x buffer (20 mM Tris-HCl, pH 8.8, 10 mM (NH_4_)_2_SO_4_, 10 mM KCl, 2 mM MgSO_4_, 0.1% Triton X-100).

### Expression and purification of ALFA Nb-Nluc

The ALFA tag nanobody (45) with an N-terminal pelB signal sequence and a C-terminal Nluc-6xHis was ordered as a gBlock (IDT) and cloned into a pBAD vector (see supplemental information for plasmid DNA sequence) by Gibson assembly (NEBuilder HiFi mix). NiCo(DE3) cells (NEB) containing the pBAD pelB-ALFA Nb-Nluc-6xHis were grown overnight in 1L of ZYM-505 media (122). After 16 hours of growth, 20 mL of 1 M NaOH was added to each culture, the protein expression was induced with the addition of 2.5 mL of 20% arabinose, and grown at 37 °C for 4 hours. Following expression the ALFA Nb-Nluc protein was purified by periplasmic protein prep as follows. The cells were pelleted by centrifugation and the cell pellet (typically ∼11,000 ODV per L expression) was resuspended in 22.5 mL ice cold TES buffer (200 mM Tris-HCl, pH 8, 0.65 mM EDTA, 0.5 M sucrose) and incubated with rocking at 4 °C for 1 hr. To this sample, 45 mL of ice cold TES/4 buffer (1:4 dilution of TES buffer in H_2_O) was added and the resulting sample was incubated with rocking at 4 °C for 30 min. The sample was centrifuged at 10k rcf fr 30 min at 4 °C and the supernatant was filtered through a bottle top filter (Thermo Scientific 0.45 μm SFCA membrane) to yield the soluble periplasmic extract. The soluble periplasmic extract was adjusted to 500 mM NaCl, 20 mM imidazole, 10 mM MgCl_2_, and 1% Triton X-100, applied to a 1 mL Ni-NTA column (Thermo Scientific HisPur Ni-NTA Resin), washed with 40 bed volumes of Ni-NTA wash buffer (50 mM Tris-HCl, pH 8, 500 mM NaCl, and 20 mM imidazole), and eluted with five sequential additions of 0.5 mL of elution buffer (Ni-NTA wash buffer supplemented with 500 mM imidazole). The resulting protein was buffer exchanged by PD-10 column (GE Healthcare) into PBS (10 mM Na_2_HPO_4_, 1.8 mM KH_2_PO_4_, pH 7.4, 137 mM NaCl, 2.7 mM KCl) mixed to a final concentration of 50% by volume with glycerol and stored at-20 °C.

### Production of ACE-tRNA PCR fragments for production of minicircles by T4 ligase-dependent *in vitro* covalent closure

PCR fragments for *in vitro* production of ACE-tRNA were obtained by amplifying the 1 kb ACE-tRNA^Arg^_UGA_ or 1 kb ACE-tRNA^Leu^_UGA_ vector (see supplemental file for plasmid DNA sequences) with either 200bp bluntF 5P (5’-/5Phos/GGCTTGATGGAGCAGTTATACTCTAAGCGC-3’) and 200 bp bluntR 5P (5’-/5Phos/CTTAGAATCACAGACTGGCATCCACCC-3’) for production of 200 bp MC or 400bp bluntF 5P (5’-/5Phos/CAGCAGAGCTTGAACTGACTAGTTGGAC-3’) and 400bp bluntR 5P (5’-/5Phos/GCTAGTTAGAAAGGCTCATCTTTGGCTAG-3’) for production of 400 bp MC. A typical PCR reaction mix contained 32.5 μL MQ-H_2_O, 10 μL 5x buffer (150 mM Tris-HCl, pH 10, 200 mM K_2_SO_4_, 5 mM (NH_4_)_2_SO_4_, 10 mM MgSO_4_, 0.5% Triton X-100), 1 μL dNTPs (10 mM each dNTP), 2.5 μL forward primer, 2.5 μL reverse primer, 1 μL plasmid template DNA (10 ng/μL), and 1 μL of the Pfu-Sso7d polymerase stock. A typical reaction was scaled up to 50 mL with the PCR reaction mix aliquoted into PCR plates at 100 μL per well. The PCR cycle utilized was 98 °C for 30 seconds, [98 °C for 10 seconds, 65 *°*C for 10 seconds, and 72 °C for 20 seconds], the previous section in brackets was repeated 30 times, a final extension cycle at 72 °C for 2 minutes was conducted, and the product was held in the thermocyclers at 12 °C. The resulting PCR product was pooled and purified by anion exchange chromatography typically at midiprep (per 50 mL PCR reaction volume) scale (Macherey-Nagel). The pooled PCR product was mixed with an equal volume of 2xEquilibration buffer (1.8 M KCl, 200 mM Tris-HCl, pH 6.3, 30% ethanol, and 1% Triton X-114) and applied to a midiprep column pre-equilibrated with 1xEquilibration buffer. After application of the crude PCR product, the midiprep column was washed with 35 mL of wash buffer 1 (1.15 M KCl, 100 mM Tris-HCl, pH 6.3, 15% ethanol, and 0.5% Triton X-114), followed by 15 mL of wash buffer 2 (1.15 M KCl, 100 mM Tris-HCl, pH 6.3, and 15% ethanol). After the wash buffer was allowed to flow through the column, the PCR product was eluted from the column with 5 mL of elution buffer (1.5 M KCl, 100 mM Tris-HCl, pH 7.0, 15% isopropanol). The eluted sample was adjusted with the addition of 3.5 mL of isopropanol, and after inverting the mixture several times, the DNA was allowed to precipitate overnight at room temperature. The sample was centrifuged in a fixed angle rotor at 20k rcf, for 45 minutes at 4 °C. The resulting DNA pellet was transferred to a microcentrifuge tube and washed 3 times with 1.5 mL of 70% ethanol, allowed to dry for 10 minutes, and dissolved in 1x T.E. buffer, pH 8.0 (Fisher Scientific). For a 200 bp minicircle assembly, a small scale 400 μL ligation reaction was composed of 299 μL MQ H_2_O, 4 μg purified PCR product, 40 μL 10x T4 DNA ligase buffer (500 mM Tris-HCl and 100 mM MgCl_2_), 40 μL ATP (10 mM stock), 4 μL DTT (freshly prepared 1 M stock), 12 μL TFAM (50 μM stock), and 1 μL T4 DNA ligase (∼400 units/μL). For a 400 bp minicircle assembly, a small scale 200 μL reaction was composed of 147 μL MQ H_2_O, 4 μg purified PCR product, 20 μL 10x T4 DNA ligase buffer (500 mM Tris-HCl and 100 mM MgCl_2_), 20 μL ATP (10 mM stock), 2 μL DTT (freshly prepared 1 M stock), 6 μL TFAM (50 μM stock), and 1 μL T4 DNA ligase (∼400 units/μL). The components were added to the reactions in the order listed and the ligation reactions were allowed to proceed overnight in the dark at ambient room temperature. Residual unreacted MC PCR product was removed by addition of T5 exonuclease (NEB, M0663S) with ∼20 μL of 10 unit/μL T5 exonuclease added per 50 mL MC ligation reaction. The reaction containing the T5 exonuclease was allowed to proceed at 37 °C for 4 hours. The intact ACE-tRNA MC DNA was purified from the rest of the reaction by anion exchange column as outlined above for purification of the PCR product.

### Production of ACE-tRNA minicircles by *in vivo* recombination

The pMC.CMV-GFP-SV40PolyA parental minicircle positive control plasmid vector (System Biosciences) was digested with XmaI/BstEII and the resulting linearized backbone was purified by agarose gel electrophoresis and gel purification kit (NEB, Monarch DNA gel extraction kit). A CmR/ccdB selection cassette was amplified from an Addgene plasmid #62176 (124) flanked by BbsI restriction sites for Golden Gate assembly of multiple ACE-tRNA copies. The pMC producer plasmids containing ACE-tRNAs were transformed into the ZYCY10P3S2T producer bacterial strain. Transformed bacterial cultures were grown for 8 hours in 2 mL starter cultures containing kanamycin before inoculating 500 mL of ZYM-505 (122) media containing kanamycin, which were grown overnight at 37 °C, 230 rpm in a baffled 2.8 L Fernbach flask. After growing overnight, the culture was supplemented with 500 mL fresh LB media, 20 mL of 1 M NaOH, and 500 μL 20% L-arabinose. The induced culture was maintained at 32 °C, 230 rpm for 5 hours.

For large scale minicircle DNA purification a maxiprep kit (Macherey-Nagel) was used with some modifications. The ODV (OD_600_ x total volume of culture) of the culture was determined (for a typical purification an ODV of 180,000 was harvested from 12 L of bacterial culture), with the cells resuspended in 0.016 mL RES-EF (containing RNase A) per ODV, lysed in an equal volume of LYS-EF, and neutralized with an equal volume of NEU-EF. After incubating on ice for 1 hour, the lysate was clarified with a combination of centrifugation, straining with cheesecloth, and bottle top filter (NucleoBond Bottle Top Filter Type 2, Macherey-Nagel). The minicircles were purified from the clarified lysate from this step as per the manufacturer instructions (NucleoBond Xtra Maxi EF, Macherey-Nagel). To remove residual parental plasmid and miniplasmid, the resulting minicircle DNA was digested overnight at 37 °C with 10 μL Esp3I (NEB, R0734S) and 20 μL T5 exonuclease (NEB, M0663S) per 500 μg minicircle in 1x cutsmart buffer (NEB). The resulting minicircle DNA was purified by midiprep anion exchange column as outlined above for the PCR product. Sequences of the minicircles resulting from recombinase-based production are reported in Supplemental Table 1.

### Production of ACE-tRNA PCR fragments for production of CEDTs by TelN-dependent *in vitro* reaction

A pUC57 Kan plasmid containing two *telRL* sites flanking a Golden Gate site was synthesized (Genscript). A CmR/ccdB selection cassette flanked by BbsI restriction sites was cloned between the *telRL* sites by Gibson assembly (NEBuilder HiFi master mix). Different length stretches of DNA containing either ACE-tRNA^Arg^_UGA_ or ACE-tRNA^Leu^_UGA_ were amplified by PCR (Q5 polymerase, NEB) and cloned by Golden Gate assembly into the pUC57 Kan 2x*telRL* plasmid (see supplemental information for plasmid DNA sequences). For the 200 bp CEDT the ACE-tRNAs flanked by *telRL* sites were cloned back-to-back sharing a single *telRL* site between cassettes (Fig. 1C) by Golden Gate assembly of gBlocks (IDT), which reduces the number of *telRL* sites that need to be cleaved per 2 individual 200 bp ACE-tRNA CEDTs from 4 sites to 3 sites. For production of ACE-tRNA CEDTs, the ACE-tRNA *telRL* plasmid DNA served as a template for PCR amplification, which was carried out as outlined above for the MC PCR. In this case the primers used were CEDTprodF (5’-GTTGTAAAACGACGGCCAGAGAATTC-3’) and CEDTprodR (5’-CAGCTATGACCATGATTACACCAAGC-3’), which anneal to the 2x *telRL* pUC57 plasmid just outside of the *telRL* sites. The CEDT PCR products were purified as outlined above for the MC PCR product. Addition of concentrated TelN directly to concentrated DNA solutions was noted to result in precipitate formation, so a dilute solution of TelN in reaction buffer was mixed with a dilute solution of ACE-tRNA *telRL* DNA. A typical 5 mL DNA mixture was composed of 100 ng/μL ACE-tRNA *telRL* DNA in 1x buffer (20 mM Tris-HCl, pH 8.8, 10 mM (NH_4_)_2_SO_4_, 10 mM KCl, 2 mM MgSO_4_, 0.1% Triton X-100). A typical 5 mL TelN mixture was composed of the appropriate amount of TelN for the number of *telRL* sites possessed by the ACE-tRNA substrate in question (assuming a TelN concentration of ∼20 units/μL, 1.4 mL of TelN was used for a 200 bp substrate, 1 mL of TelN was used for a 400 bp substrate, 0.6 mL of TelN was used for a 900 bp substrate) in 1x buffer. The dilute DNA and TelN solutions were mixed, gently inverted several times, and the reaction was allowed to proceed at 30 °C for 2 hours. The resulting digested DNA mixture was purified by anion exchange maxiprep column as outlined above for purification of PCR product, the residual undigested fragments and cleaved open ended fragments were removed by T5 exonuclease digest, and the final product was purified by anion exchange midiprep column as outlined above for the PCR product.

**Figure 1.**
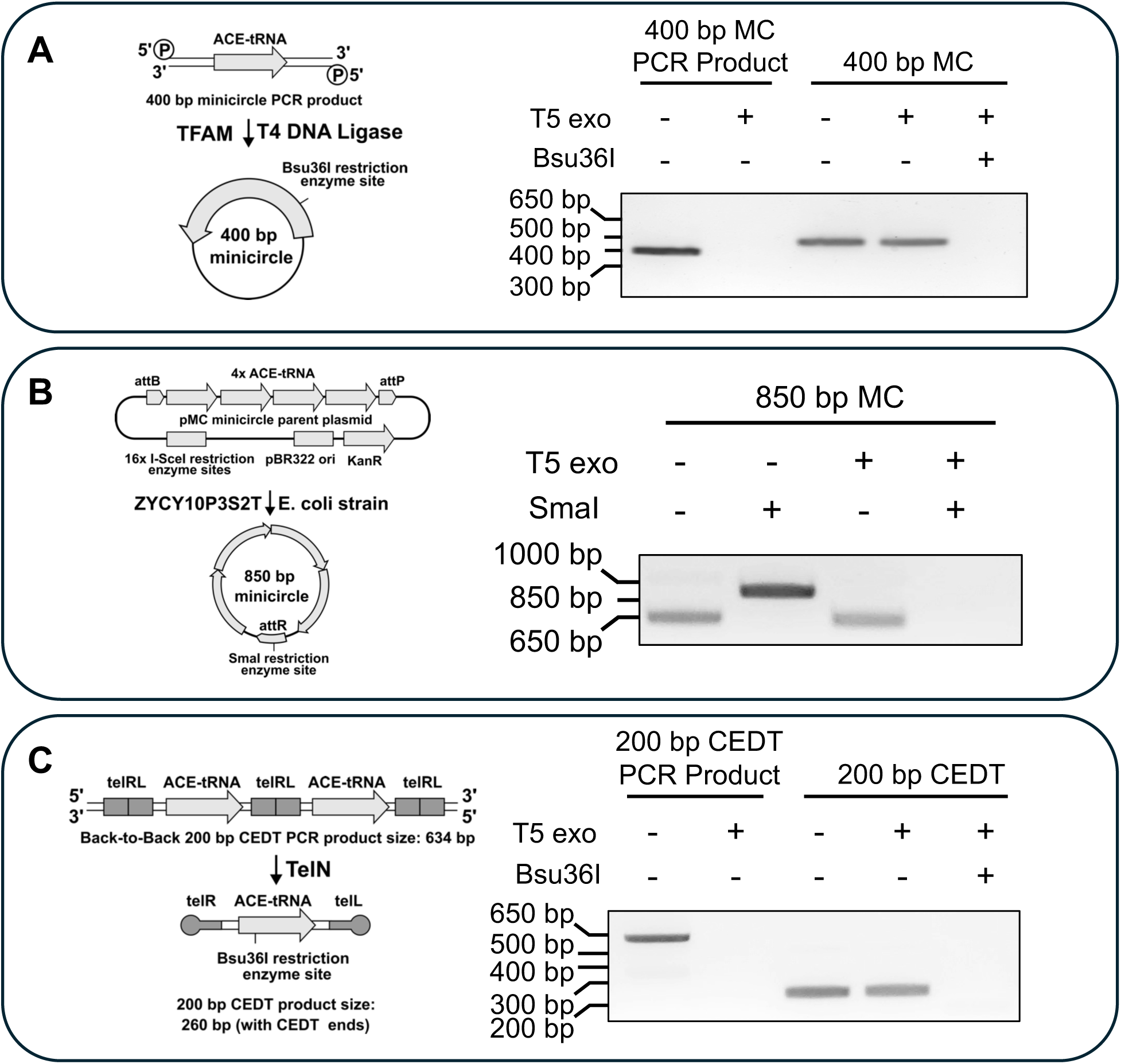
ACE-tRNAs can be encoded in minimal DNA vectors (minivectors). (**A**) Circular dsDNA minivectors (minicircles) containing a single copy of an ACE-tRNA expression cassette were produced from blunt end ligation of ACE-tRNA-containing PCR products. Primers containing a 5’-phosphate were utilized to allow for ligation with T4 DNA ligase and the nonspecific DNA-binding/bending activity of the protein TFAM was used to ensure efficient circularization of minicircle product (36). Production of T5 exonuclease-resistant product in the absence of specific endonuclease activity (Bsu36I) indicates the covalent closure of the minicircles. (**B**) Circular dsDNA minivectors containing multiple copies of an ACE-tRNA expression cassette were produced using a commercially available minivector production system, which allows for plasmid vector backbone removal based on the activity of PhiC31 integrase and I-SceI homing endonuclease in the ZYCY10P3S2T *E. coli* strain (described in (38)). (**C**) Linear dsDNA minivectors with covalently closed DNA ends (CEDTs), containing either a single or multiple ACE-tRNA expression cassette(s), were produced from PCR products containing 54 bp telRL sites, which are recognized by the cleaving/joining enzyme TelN (41). Production of T5 exonuclease-resistant product in the absence of specific endonuclease activity (Bsu36I) indicates the covalent closure of the CEDT ends.

**Figure 2.**
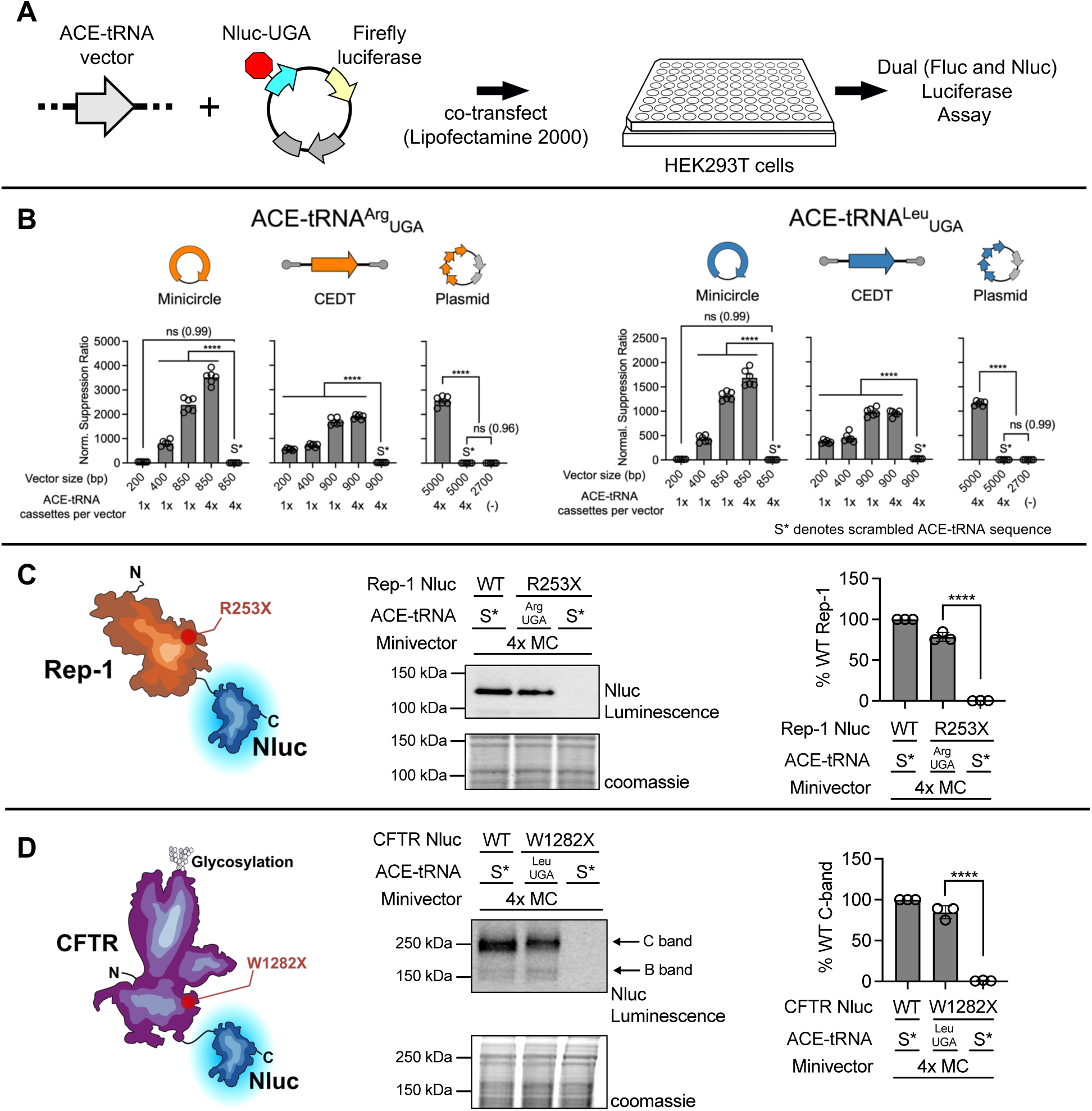
ACE-tRNAs encoded as DNA minivectors can rescue expression of PTC-containing proteins. (**A**) Function of ACE-tRNA minivectors was screened by co-transfecting the respective minivectors along with a plasmid containing a PTC-interrupted nanoluciferase (Nluc-UGA) expression cassette and a firefly luciferase (Fluc) expression cassette, which serves as a transfection control (16). (**B**) Results of dual luciferase assays (n = 6) for each minivector. Minivectors containing either ACE-tRNA^Arg^_UGA_ or ACE-tRNA^Leu^_UGA_, with the size and DNA structure noted, were tested. The normalized suppression ratio shown here is calculated from the equation (PTC-NanoLuciferase luminescence [+ACE-tRNA]/Firefly luminescence)/(PTC-Nanoluciferase luminescence [no ACE-tRNA]/Firefly luminescence). S* denotes a minivector or plasmid containing a scrambled ACE-tRNA, which serves as a negative control. (**C**) 850 bp 4xACE-tRNA^Arg^_UGA_ minicircle demonstrates significant rescue of Rep-1 p.R253X, the most common PTC that causes choroideremia. A Rep-1 construct containing the p.R253X variant and a C-terminal Nluc fusion tag was co-transfected (n = 3) along with a 4xACE-tRNA^Arg^_UGA_ minicircle in the HEK293T cell line. SDS-PAGE in-gel Nluc luminescence band densitometry was used to assay for production of full-length Rep-1 with subsequent Coomassie staining serving as a loading control. (**D**) 850 bp 4xACE-tRNA^Leu^_UGA_ minicircle demonstrates significant rescue of CFTR p.W1282X, a common PTC that causes cystic fibrosis. As shown above, a CFTR construct containing the p.W1282X variant and a C-terminal Nluc fusion tag was co-transfected (n = 3) along with a 4xACE-tRNA^Leu^_UGA_ minicircle in the HEK293T cell line. SDS-PAGE in-gel Nluc luminescence band densitometry was used to assay for production of full-length fully glycosylated (C-band) CFTR with subsequent Coomassie staining serving as a loading control. Data are presented as the mean ± SEM with significance determined by one-way ANOVA and Tukey’s post-hoc test, where **** *p* < 0.0001, and ns denotes not significantly different (*p* value shown for each ns).

**Figure 3.**
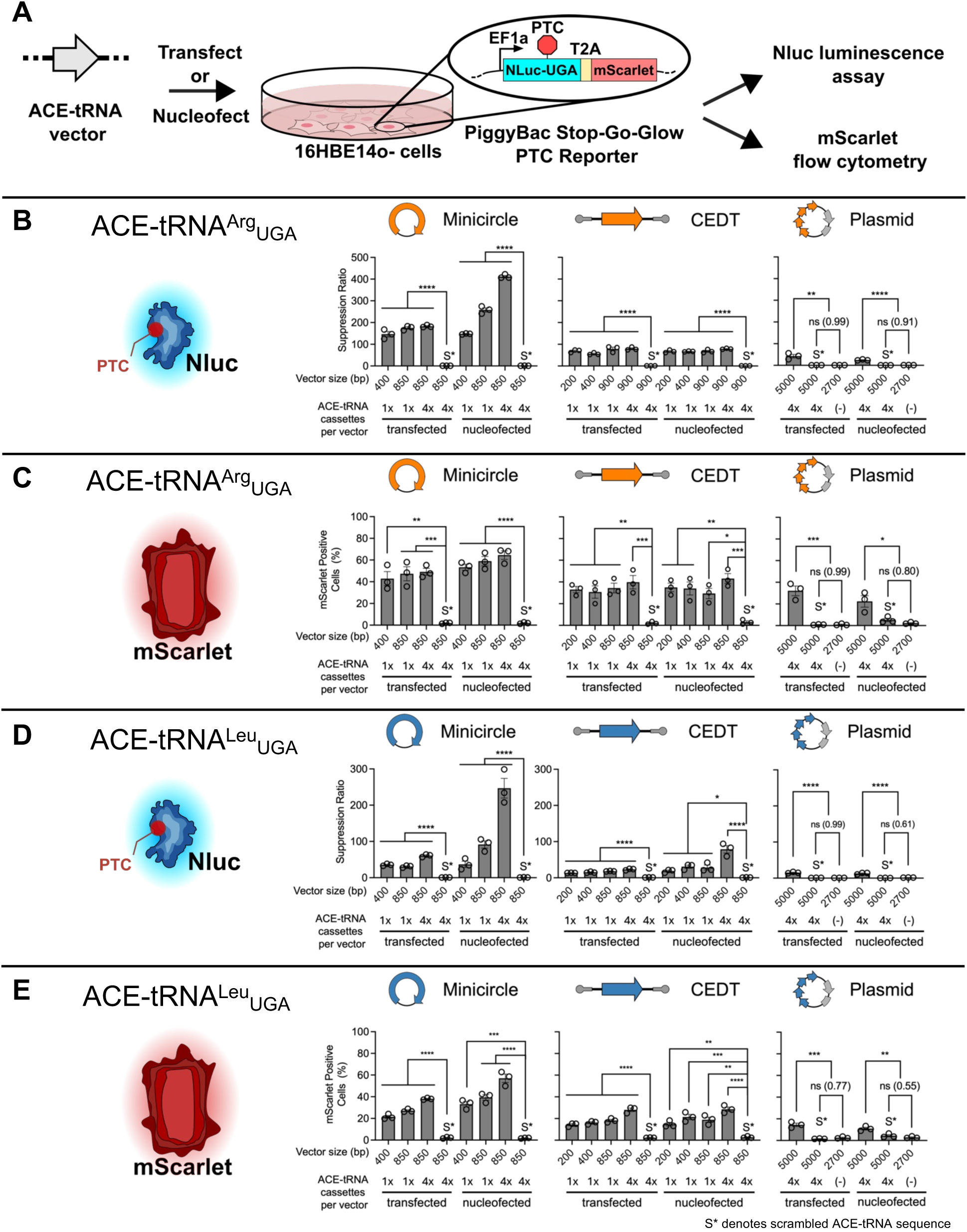
ACE-tRNAs encoded as DNA minivectors can support efficient PTC rescue when delivered by lipofectamine mediated transfection and by nucleofection to human bronchial epithelial cells. (**A**) The Stop-Go-Glow (SGG) PTC reporter containing a PTC-interrupted nanoluciferase (Nluc-UGA) and a downstream mScarlet separated by a 2A element, was stably integrated into 16HBE14o-cells using the piggyBac transposon system as described previously (15). The PTC suppression activities of ACE-tRNA^Arg^_UGA_ and ACE-tRNA^Leu^_UGA_ minivectors were determined by NanoLuciferase luminescence via plate reader assay (**B** and **D**, respectively) and mScarlet flow cytometry (**C** and **E**, respectively). S* denotes a minivector or plasmid containing a scrambled ACE-tRNA, which serves as a negative control. The suppression ratio shown here is calculated from the equation (PTC-NanoLuciferase luminescence [+ACE-tRNA])/(PTC-Nanoluciferase luminescence [no ACE-tRNA]). Data are presented as the mean ± SEM (n = 3) with significance determined by one-way ANOVA and Tukey’s post-hoc test, where * *p* < 0.05, ** *p* < 0.01, *** *p* < 0.001, **** *p* < 0.0001, and ns denotes not significantly different with *p* values shown for these cases.

### Production of supercoiled ACE-tRNA minicircles

Supercoiled 200 bp and 400 bp ACE-tRNA minicircles were produced by adapting previously outlined methods (125). Open circle minicircles were digested with the nicking endonuclease Nt.AlwI, supercoiling was induced with application of 6 μg/mL ethidium bromide, and the nicked DNA was sealed with T4 DNA ligase, with residual nicked DNA removed with T5 exonuclease. Supercoiled minicircles were resolved by agarose gel electrophoresis and purified by gel purification kit (NEB, Monarch DNA gel extraction kit).

### Transfection of ACE-tRNA minivectors and pDNA

HEK293T cells were cultured and transfected using Lipofectamine 2000 as previously described (16). ACE-tRNA minivectors and pDNA were transfected at equal mass with the pNanoRePorter plasmid, with the PTC suppression assay conducted using the Nano-Glo Dual-Luciferase Reporter Assay System (Promega) as previously described (16) and as shown in the supplemental information (Fig. S5A-C).

### Selection of the 16HBE14o-Stop-Go-Glow cell line

The Stop-Go-Glow (SGG) reporter used here was composed of an EF1a-promoter driven cassette expressing a PTC-interrupted Nluc reporter (12), a T2A ribosomal shutter element, and a downstream mScarlet-I (Fig. S5D), which along with a puromycin resistance cassette was cloned between *piggyBac* ITRs (15) for selection of a 16HBE14o-stable cell line as described previously (22).

### Transfection of the 16HBE14o-Stop-Go-Glow cell line for Nluc assay in 96-well format

The 16HBE14o-SGG stable cell line was transfected (Lipofectamine LTX, Thermo Fisher Scientific) with ACE-tRNA minivectors and pDNA. For transfection of the minivectors for the Nluc assay, the 16HBE14o-SGG cells were trypsinized (TrypLE Express, Gibco), the cells were counted according to manufacturer instructions (Countess 3 Automated Cell Counter, Thermo Fisher Scientific), and diluted to 3 x 10^5^ cells. To each well of a fibronectin coated (as described previously (15)) cell-culture treated, black, flat-bottom 96-well plate (Greiner Bio-One), 100 μL of the diluted cells were added and the plate was placed in a 37 °C/5% CO_2_ incubator. The minivector or pDNA was diluted to 200 ng/μL and 2 μL was used for transforming 4 wells of the 96-well plate. PLUS reagent (Lipofectamine LTX kit) was mixed with OptiMEM (Gibco) at a ratio of 0.4 μL PLUS reagent per 20 μL OptiMEM, while the LTX reagent (Lipofectamine LTX kit) was mixed with OptiMEM at a ratio of 0.8 μL LTX per 20 μL OptiMEM. Each DNA sample had 20 μL of the PLUS/OptiMEM mix added and was thoroughly mixed by pipetting several times, this sample then had 20 μL of the LTX/OptiMEM mix added and was thoroughly mixed by pipetting several times. The resulting DNA mixture was incubated for ∼7.5 minutes at room temperature, before 10 μL of the mixture was added to each respective well of cells. The transfected cells were replaced in the 37 °C/5% CO_2_ incubator for 24 hr before PTC-suppression was assayed using the Nano-Glo Luciferase Assay System (Promega) (Fig. S5D-E). To carry out the Nluc assay, the cell media was aspirated, 15 μL of DPBS (Gibco, Cat. # 14190144) was added to each well, the Nano-Glo Luciferase Assay Substrate (furimazine) was diluted at a 1:50 ratio in Nano-Glo Luciferase Assay Buffer, 15 μL of the diluted substrate was added to each well of cells, the plate was incubated shaking at 600 rpm for 3 minutes, and the Nluc luminescence was measured by a Synergy2 multi-mode microplate reader (Biotek Instruments). The suppression ratio is calculated from the equation ([Nluc luminescence value (+ACE-tRNA MV or pDNA well)]/ [Nluc luminescence value (+scrambled negative control ACE-tRNA MV or pDNA well)]) (Fig. S5F).

### Nucleofection of the 16HBE14o-Stop-Go-Glow cell line for Nluc assay in 96-well format

To each well of a fibronectin coated (as described previously (15)) cell-culture treated, black, flat-bottom 96-well plate (Greiner Bio-One), 100 μL of the cell culture media was added and the plate was placed in a 37 °C/5% CO_2_ incubator. For each sample being nucleofected, 2 x 10^5^ cells were centrifuged at 90 rcf for 10 min, resuspended in 20 μL of SG 4D-Nucleofector X solution containing Supplement 1 (Amaxa SG cell line 4D-Nucleofector X Kit S, Lonza Bioscience), and 1 μg of each minivector or pDNA was added to the solution. A 20 μL total volume of the cell/DNA mixture was transferred to each well of a Nucleocuvette Strip. The cells were nucleofected by placing the Nucleocuvette Strip into the 4D-Nucleofector X Unit and running the CM-137 Nucleofector program from the 4D-Nucleofector Core Unit. The cells were incubated at room temperature for 10 minutes before being resuspended in 80 μL of pre-warmed cell culture media. The 96-well plate containing pre-warmed media had 15 μL of the nucleofection mix added to each well and was maintained in a 37 °C/5% CO_2_ incubator for 24 hours. The Nluc assay was conducted as described above for the Lipofectamine LTX transfection.

### Transfection of the 16HBE14o-Stop-Go-Glow cell line for flow cytometry in 12-well format

The 16HBE14o-SGG stable cell line was transfected (Lipofectamine LTX, Thermo Fisher Scientific) with ACE-tRNA minivectors and pDNA as described above in a 96-well format but with the following modifications for 12-well format. The trypsinized cells were diluted to 3 x 10^5^ cells/mL and 1 mL of cell suspension was added to each well of the 12-well plate. The DNA was diluted to 200 ng/μL and 5 μL was used to transfect each well of cells. PLUS reagent (Lipofectamine LTX kit) was mixed with OptiMEM (Gibco) at a ratio of 1 μL PLUS reagent per 50 μL OptiMEM, while the LTX reagent (Lipofectamine LTX kit) was mixed with OptiMEM at a ratio of 2 μL LTX per 50 μL OptiMEM. Each DNA sample had 50 μL of the PLUS/OptiMEM mix added and was thoroughly mixed by pipetting several times, this sample then had 50 μL of the LTX/OptiMEM mix added and was thoroughly mixed by pipetting several times. The resulting DNA mixture was incubated for ∼7.5 minutes at room temperature, before 100 μL of the mixture was added to each respective well of cells. The cells were then replaced in a 37 °C/5% CO_2_ incubator for 24 hours before conducting the flow cytometry assay.

### Nucleofection of the 16HBE14o-Stop-Go-Glow cell line for flow cytometry in 12-well format

Nucleofection of the ACE-tRNA minivector and pDNA was conducted as outlined above for the Nluc 96-well format with the following modifications. After nucleofection the 20 μL of nucleofected cells were resuspended in 80 μL of pre-warmed cell culture media. A total of 100 μL of the resuspended cells were transferred to each well of the 12-well plate, which contained 1 mL of pre-warmed cell culture media. The cells were then replaced in a 37 °C/5% CO_2_ incubator for 24 hours before conducting the flow cytometry assay.

### Preparation of SGG cell samples for flow cytometry and flow cytometry conditions

After cells were treated with ACE-tRNA minivectors and pDNA as described above, flow cytometry was conducted using the SGG mScarlet-I as a fluorescent marker. To prepare the cell samples, the media was aspirated, 300 μL of TrypLE Express (Gibco) was added to each well containing cells, the plate was placed in a 37 °C/5% CO_2_ incubator for 10 minutes, and the reaction in each well was quenched with 1 mL of FACS buffer (DPBS (Gibco, Cat. # 14190144) containing 0.3% cell culture grade BSA (Gibco, Cat. # 15260-037) and 1 mM EDTA). The trypsinized cells were pipetted up-and-down several times to dislodge the cells before being transferred to flow cytometry tubes containing 2 mL of FACS buffer. The cells were centrifuged at 200 rcf for 5 min, the FACS buffer was aspirated, and the cells were resuspended in 250 μL of fresh FACS buffer. The cell samples were run on a BD LSR Fortessa flow cytometer (BD Biosciences) flow cytometer collecting over 20,000 events. mScarlet-I was excited with a yellow-green (561 nm, 50 mW) laser and detected between 600 and 620 nm. Data were analyzed with FCS Express 7 (De Novo Software) with gating for single cells accomplished using forward and side scatter area, height, and width parameters. A gate for mScarlet-I positive cells in samples was established such that ∼1% of events were positive for cells transfected or nucleofected respectively with pUC57 pDNA without an insert. Independent mScarlet-I fluorescence gates were established for samples that were either transfected or nucleofected as the level of background autofluorescence was noted to be different for each method of DNA delivery. The percent mScarlet-I positive cells and median fluorescence intensity (MFI) of the positive population for each ACE-tRNA minivector and pDNA treated cell sample was calculated based on the gate statistical analysis in FCS Express 7.

### PTC-Rep1-Nluc and PTC-CFTR-Nluc expression and in-gel luminescence

CFTR and Rep-1 cDNA with a C-terminal Nluc fusion was cloned via Gibson assembly downstream of a short ubiquitin C (UbC) promoter in pUC57 mini. PTCs were introduced into the Rep-1 and CFTR cDNA construct using PCR based site-directed mutagenesis. The Rep-1 and CFTR expression plasmids were co-transfected along with ACE-tRNA minivector or pDNA into HEK293T cells using Lipofectamine LTX (Thermo Fisher). For all transfections 750 ng of Rep-1-Nluc or CFTR-Nluc plasmid was used, while 750 ng of ACE-tRNA minivector or pDNA was used in the DNA mix. HEK293T cells were counted and diluted to 6 x 10^5^ cells per mL with 1 mL added per well of a 6-well plate. For each well, the DNA mix had 150 μL of OptiMEM containing 3 μL of PLUS reagent (Thermo Fisher) added to and mixed with the DNA, followed by 150 μL of OptiMEM containing 9 μL of LTX reagent (Thermo Fisher). The DNA/Lipofectamine mixture was incubated for ∼7.5 minutes at room temperature, before 250 μL of the mixture was delivered dropwise to each well of cells. All transfections were performed in triplicate. The preparation of samples and in-gel luminescence assay was performed as described previously (16).

### Comparison of PTC suppression efficiency of 200 bp CEDT ACE-tRNA^Arg^_UGA_ and optimized (Opt) ACE-tRNA^Arg^_UGA_

200 bp ACE-tRNA^Arg^_UGA_ and OptACE-tRNA^Arg^_UGA_ CEDTs were produced by PCR and TelN digest as described above. The resulting CEDTs were diluted to both 150 ng/μL and 100 ng/μL and mixed at equal volume with pNanoRePorter plasmid at a concentration of 50 ng/μL. A 2-fold serial dilution was then conducted with each of the starting DNA stocks to provide the DNA amounts shown (Fig. 4B). The DNA mix for the serial dilution was composed of pNanoRePorter plasmid at 50 ng/μL mixed at equal volume with 150 ng/μL pUC57 pDNA without insert used as inert carrier DNA. Transfections into HEK293T and Nano-Glo Dual-Luciferase reporter assays were carried out as indicated above (section labeled “Transfection of ACE-tRNA minivectors and pDNA”).

**Figure 4.**
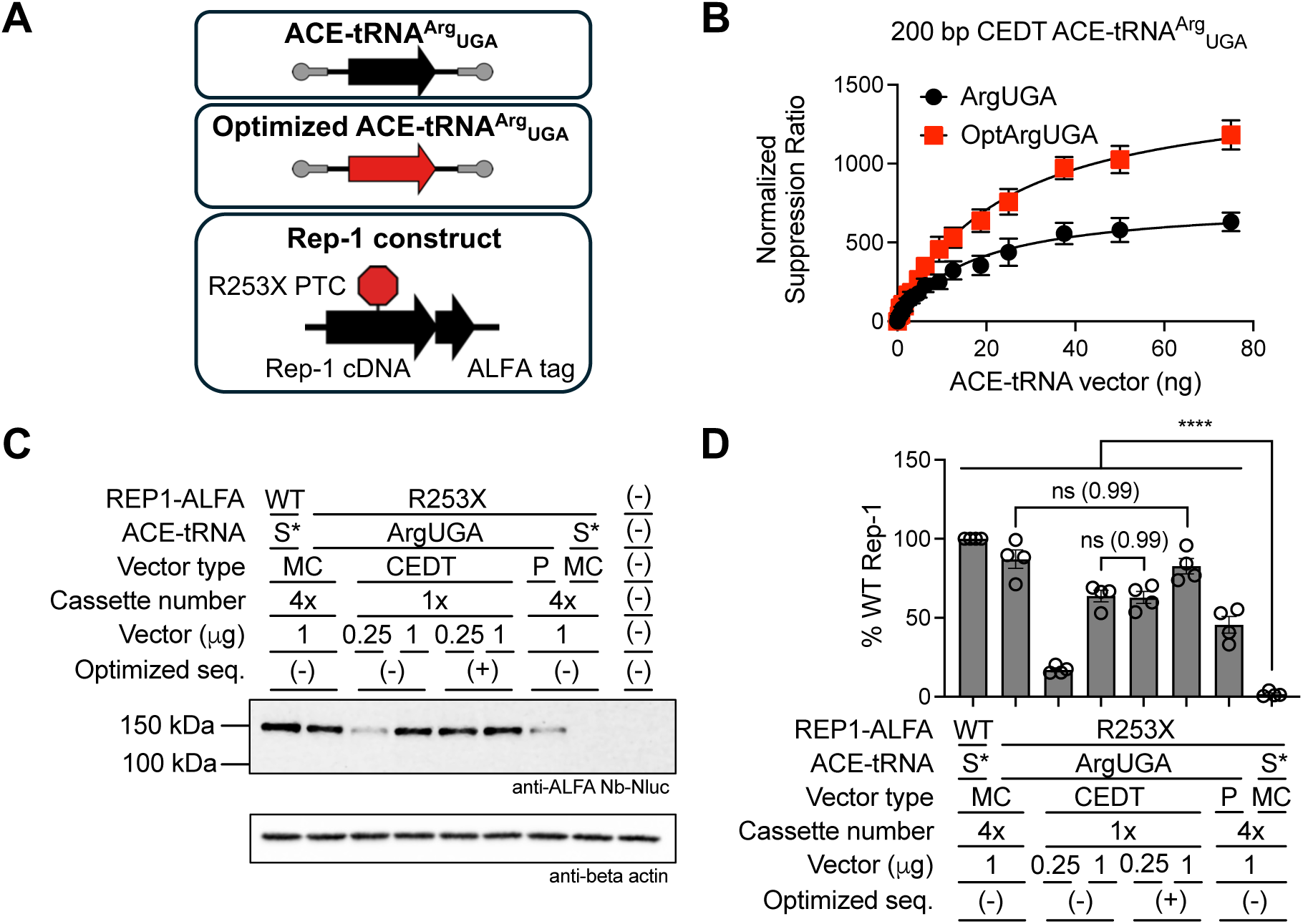
Optimized ACE-tRNAs encoded as DNA minivectors can rescue expression of Rep-1 p.R253X in a DNA concentration-dependent fashion. (**A**) Constructs used for this experiment. 200 bp CEDTs containing either ACE-tRNA^Arg^_UGA_ or Optimized ACE-tRNA^Arg^_UGA_ expression cassettes (16) and a construct containing a UbC-promoter-driven Rep-1-R253X-ALFA tag expression cassette. (**B**) Various concentrations of each of the two CEDTs shown in panel A were transfected along with the dual-luciferase PTC reporter plasmid (as shown in Fig. 2A) in HEK293T cells. The data shown is the average of 9 independent transfections with the error bars depicting the SEM and fits according to the equation outlined in (Fig. S11). (**C**) Representative western blot image showing Rep-1-253X rescue following co-transfection (n = 4). All CEDTs used in this experiment were 200 bp in size, while the MC were 850 bp in size. (**D**) Quantified band densitometry data for the Rep-1-ALFA tag western blots with data shown as the percent of WT band intensity normalized to the beta-actin loading control. Abbreviations: Minicircles, MC; Closed End DNA Threads, CEDT; Plasmid DNA, P; Scrambled ACE-tRNA Control, S*. Data are presented as the mean ± SEM with significance determined by one-way ANOVA and Tukey’s post-hoc test, where **** *p* < 0.0001, and ns denotes not significantly different (*p* value shown for each ns).

### Rep-1 western blot

The *Rep-1* cDNA with C-terminal ALFA tag was synthesized as a gBlock (IDT) and cloned into a pUC57 mini vector containing a short UbC promoter and SV40 polyA signal by Gibson assembly (NEBuilder HiFi master mix). The p.R253X variant (CGA > TGA) was inserted by site-directed mutagenesis with primers containing the desired DNA mutation (Q5 master mix, NEB). All DNA vectors were diluted to 100 ng/μL, with a typical DNA transfection mixture composed of 30 μL Rep-1 pDNA mixed with 10 μL ACE-tRNA minivector or pDNA. In cases where less ACE-tRNA minivector was transfected, the total DNA mass was balanced with pUC57 pDNA without insert, which was used as inert carrier DNA. The HEK293T cells were diluted to 6 x 10^5^ cells/mL and 1 mL was added to each well of a 6-well plate. The DNA had 160 μL of OptiMEM added, the diluted DNA then had 200 μL of OptiMEM containing Lipofectamine 2000 (Thermo Fisher Scientific) added (mixed at a ratio of 5.5 μL Lipofectamine 2000 per 194.5 μL OptiMEM). The resulting transfection mixture was thoroughly mixed by pipetting, incubated at room temperature for 10 minutes, with 150 μL added dropwise to each well of cells. The transfected HEK293T cells were maintained in a 37 °C/5% CO_2_ incubator for 24 hours before the media was aspirated, the cells washed with DPBS, and lysed in RIPA buffer (Invitrogen) containing protease inhibitors (Medchem Express). Insoluble cell debris was removed by centrifugation at 20k rcf for 30 minutes at 4 °C, total protein was quantified by BCA assay (Pierce), and 10 μg of protein was resolved on a 10-20% Novex WedgeWell SDS-PAGE gel (Invitrogen). The protein was transferred from the gel to 0.45 μm Immobilon-FL PVDF membrane (Merck Millipore) in Towbin’s transfer buffer (25 mM Tris, 192 mM glycine, 20% methanol) at 50 V, 4 °C, for 1 hour. After transfer, the membrane was washed once in MQ-H_2_O and blocked for 1 hour with rocking in EveryBlot Blocking Buffer (Biorad). The membrane was then probed with ALFA Nb-Nluc (purification outlined above) at 1:1k and mouse (monoclonal AC-15) anti-beta-actin (Fisher, AM4302) at 1:50k in buffer composed of equal volumes of EveryBlot Blocking Buffer and low salt TBST (50 mM Tris-HCl, pH 7.4, 7.5 mM NaCl, 0.1% Tween 20). The membrane was incubated for 1 hour at room temperature with rocking in the buffer containing the antibodies. The primary antibody solution was then removed, and the membrane was washed 4 times for 15 minutes each in low salt TBST. For detection of the anti-beta-actin the membrane was then probed with IRDye 800CW goat anti-mouse IgG (H+L) antibody (Licor Bio) at 1:10k in buffer composed of equal volumes of EveryBlot Blocking Buffer and low salt TBST rocking at room temperature for 1 hour. The secondary antibody solution was then removed, and the membrane was washed 4 times for 15 minutes each in low salt TBST. The membrane was incubated for 5 minutes with rocking in DPBS supplemented with 1 mM DTT and 1:500 Nano-Glo substrate (Promega). The resulting luminescence was imaged on a ChemiDoc Imaging System (Biorad) with an exposure time of 1400 sec typically used.

Quantification of Rep-1-ALFA and beta-actin band intensity was determined using Image Lab software version 6.1.0 (Biorad).

### Nucleofection of ACE-tRNA minivectors and pDNA into 16HBEge W1282X cells

ACE-tRNA minivectors and pDNA were delivered to 16HBEge W1282X cells by nucleofection for assessment of PTC-containing *CFTR* mRNA and protein rescue. 16HBEge W1282X cells were trypsinized and counted as outlined above. For each nucleofection 1.5 x 10^7^ cells were pelleted by centrifugation at 90 rcf for 10 minutes. The cells were then gently resuspended in 100 μL of SG 4D-Nucleofector X solution containing Supplement 1 (Amaxa SG cell line 4D-Nucleofector X Kit L, Lonza Bioscience). Each tube of resuspended cells had 10 μg (∼1% of the total volume of the solution) of ACE-tRNA minivector or pDNA added and the solution was mixed gently twice. 100 μL of the resuspended cell and DNA solution was transferred to the Nucleocuvette, which was placed into the 4D-Nucleofector X Unit with the CM-137 protocol subsequently run on the 4D-Nucleofector Core Unit. The cells were incubated at room temperature for 10 minutes before being resuspended in 550 μL of pre-warmed cell culture media. The nucleofected cells were then transferred to one well of a fibronectin coated 6-well plate containing 1 mL of pre-warmed cell media and maintained in a 37 °C/5% CO_2_ incubator. After 16 hours the media was aspirated and replaced with fresh media. After 48 hours the cells were either trypsinized for mRNA isolation or lysed in RIPA buffer (Invitrogen) containing protease inhibitors (Medchem Express) for western blot analysis. The nucleofection was performed three independent times.

### Quantification of *CFTR* PTC-containing mRNA by RT-qPCR from 16HBEge W1282X cells treated with ACE-tRNA minivectors and pDNA

ACE-tRNA minivectors and pDNA were delivered to 16HBEge W1282X cells by nucleofection as described above. Trypsinized cells were pelleted by centrifugation and stored at-80 °C before total RNA was isolated (Monarch Total RNA Miniprep Kit, NEB). PTC-containing *CFTR* mRNA transcript from three independent nucleofections was quantified by RT-qPCR, which was conducted as described previously (15).

### Quantification of *CFTR* PTC-containing protein from 16HBEge W1282X cells treated with ACE-tRNA minivectors and pDNA

ACE-tRNA minivectors and pDNA were delivered to 16HBEge W1282X cells by nucleofection as described above. Cells for western blot analysis were washed with DPBS, and lysed in RIPA buffer (Invitrogen) containing protease inhibitors (Medchem Express). Insoluble cell debris was removed by centrifugation at 20k rcf for 30 minutes at 4 °C, total protein was quantified by BCA assay (Pierce), and 10 or 30 μg of total protein was resolved on 4-12% Novex WedgeWell SDS-PAGE gels (Invitrogen). After running the gel was washed once in MQ-H_2_O and equilibrated in Towbin’s transfer buffer (25 mM Tris, 192 mM glycine, 10% methanol) for 15 minutes at room temperature with rocking. The protein was transferred from the gel to 0.22 μm supported nitrocellulose membrane (GVS North America, Ref 1212721) in Towbin’s transfer buffer (25 mM Tris, 192 mM glycine, 10% methanol) at 30 V, 0.09 A max per gel, 4 °C, for 16 hours. After transfer, the membrane was washed once in MQ-H_2_O and blocked for 1 hour with rocking in EveryBlot Blocking Buffer (Biorad). The membrane was then probed with anti-CFTR antibody (University of North Carolina CFTR antibody 596) at 1:5k and rabbit (monoclonal ST0533) anti-sodium potassium ATPase antibody (Thermo Fisher, MA5-32184) at 1:5k in buffer composed of equal volumes of EveryBlot Blocking Buffer and low salt TBST (50 mM Tris-HCl, pH 7.4, 7.5 mM NaCl, 0.1% Tween 20). The membrane was incubated for 16 hours at 4 °C with rocking in the buffer containing the antibodies. The primary antibody solution was then removed, and the membrane was washed 4 times for 15 minutes each in low salt TBST. For detection of the anti-CFTR the membrane was then probed with stabilized peroxidase conjugated goat anti-mouse (H+L) antibody (Invitrogen, 32430) at 1:5k and for detection of the anti-sodium potassium ATPase the membrane was simultaneously probed with IRDye 680RD donkey anti-rabbit (Licor Bio) at 1:10k in buffer composed of equal volumes of EveryBlot Blocking Buffer and low salt TBST rocking at room temperature for 1 hour. The secondary antibody solution was then removed, and the membrane was washed 4 times for 15 minutes each in low salt TBST. To image the anti-CFTR secondary antibody with conjugated peroxidase the membrane was incubated in ECL substrate (Supersignal West Femto Maximum Sensitivity, Thermo Scientific) for three minutes before being imaged on a ChemiDoc Imaging System (Biorad). The anti-sodium potassium ATPase secondary antibody conjugated with IRDye680 was subsequently imaged on the ChemiDoc Imaging System (Biorad). Quantification of CFTR and sodium potassium ATPase band intensity was determined using Image Lab software version 6.1.0 (Biorad).

### Treatment of 16HBE14o-cells with the anti-mitotic agent aphidicolin for the determination of ACE-tRNA minivector and pDNA function in mitotically arrested cells

16HBE14o-cells containing the genomically encoded SGG PTC reporter were trypsinized, counted, and plated at 2.5 x 10^5^ cells/mL (+aphidicolin) or 1 x 10^5^ cells/mL (+vehicle), 100 μL per well in 96-well format as described above. The different cell concentrations were used to account for arrested cell division in the cells treated with aphidicolin. Cells treated with aphidicolin were maintained in complete cell media containing 5 μg/mL aphidicolin (MilliporeSigma, 1782731MG) for 16 hours before transfection, while control cells were maintained in complete cell media containing vehicle (DMSO). The cells were transfected with ACE-tRNA minivectors or pDNA using Lipofectamine 2000 as previously described (16). After 24 hours post-transfection, the Nluc-PTC assay was conducted as described above (section labeled “Transfection of the 16HBE14o-Stop-Go-Glow cell line for Nluc assay in 96-well format”).

### Determination of cGAMP production in HeLa cells treated with ACE-tRNA minivectors or pDNA by competitive ELISA

HeLa cells (ATCC) were cultured in Dulbecco’s Modified Eagle Medium (DMEM) supplemented with 10% FBS, 1% Penicillin-Streptomycin, and 2 mM L-glutamine (Thermo Fisher). Cells were seeded in a 6-well plate at 2.5 x 10^5^ cells/mL, with 2 mL of cell solution added to each well and maintained in a 37 °C/5% CO_2_ incubator for 24 hours. At a timepoint of 30-60 min before transfection, the media was aspirated and exchanged for 1 mL of fresh cell culture media. For transfection of ACE-tRNA minivector or pDNA, 1.5 μg of DNA was diluted in 60 μL of OptiMEM, with 60 μL of Lipofectamine 2000/OptiMEM (3 μL Lipofectamine 2000 per 57 μL OptiMEM) mix added to each DNA solution. The Lipofectamine 2000/DNA mix was incubated at room temperature for 5 min, before 120 μL was added dropwise to each well of cells. Three independent transfections were performed for each ACE-tRNA minivector or pDNA sample. The transfected cells were replaced in the 37 °C/5% CO_2_ incubator for 6 hours, before the media was aspirated, the cells washed with DPBS, and the cells lifted by cell scraper in 200 μL lysis buffer (20 mM HEPES, pH 7.2, 150 mM NaCl, 10 mM EDTA) before being transferred to microcentrifuge tubes. The cells were lysed by sonication, heated to 95 °C for 5 min, cooled on ice for 10 min, and the insoluble cell debris was pelleted by centrifugation at 20k rcf for 30 min. The total protein in the supernatant was quantified by BCA assay (Pierce), and the 2’-5’/3’-5’ cyclic-GMP-AMP (cGAMP) levels were assessed by competitive ELISA according to manufacturer instructions (Invitrogen, Ref EIAGAMP).

### Determination of ACE-tRNA minivector or pDNA stability in human

Pooled human serum off the clot, female and male mix (Innovative Research, ISERFM100ML), was further clarified by centrifugation at 20k rcf, 4 °C, for 30 minutes, filtered through a bottle top filter (Thermo Scientific 0.45 μm SFCA membrane), and then aliquoted and stored at-80 °C. For stability assays 2 μg of each ACE-tRNA minivector or pDNA was incubated in 200 μL of the human serum. At the timepoint indicated (Fig. 6C) the serum nucleases were quenched with the addition of 4 μL of 20 mg/mL proteinase k (NEB), and 7 μL 30% SDS, before being incubated at 60 °C for 2 hours. The 0 timepoint had the serum pre-treated with proteinase k and SDS. The samples were then extracted with 200 μL phenol/chloroform/isoamyl alcohol (25:24:1), saturated with pH 8 100 mM Tris-EDTA (Thermo Scientific). To aid with DNA extraction, 200 μL of 1x TE was added, the sample were inverted thoroughly, and the mixture was centrifuged at 20k rcf, for 10 min, at 4 °C. From the aqueous phase, 300 μL was transferred to a new microcentrifuge tube, and 400 μL of phenol/chloroform/isoamyl alcohol was added. The sample was inverted thoroughly and centrifuged as before. From this aqueous phase, 250 μL was transferred to a new microcentrifuge tube. To precipitate the DNA, 500 μL ice cold ethanol, 75 μL 3 M sodium acetate (pH 5.2), and 5 μL glycogen (20 mg/mL, Thermo Fisher) was added to the DNA, mixed thoroughly by inversion, and held at-20 °C overnight. The samples were centrifuged at 20k rcf, for 30 min, at 4 °C, the supernatant was removed, the pellets were washed with 70% ethanol, dried for 10 min, and resuspended in 20 μL TE. Each sample had 5 μL loaded on an agarose gel (2% for minivectors, 0.5% for pDNA) with DNA visualized by subsequent ethidium bromide staining and imaging with a ChemiDoc Imaging System (Biorad). Quantification of ACE-tRNA minivector and pDNA band intensity was determined using Image Lab software version 6.1.0 (Biorad). The data was fit to a one phase decay using least squares (GraphPad Prism 10.5.0).

## RESULTS

### Production of ACE-tRNA minivectors

Production of ACE-tRNA minivectors followed several previously established approaches with some modifications. Variable size, single-copy ACE-tRNA minicircles were produced by PCR rather than solid-phase oligonucleotide synthesis (36). This PCR approach allows for a range of minivector sizes to be produced from the same template depending on the primers used for amplification. Primers containing 5’ phosphates were used for PCR amplification, as this allows for blunt end ligation saving further enzymatic processing and purification steps.

Depending on the sequence of the DNA, sequences shorter than 1000 bp are increasingly difficult to bend into a circle due to the inherent DNA rigidity (37). As such, a ligase-mediated, bending protein-assisted circularization reaction was utilized to produce small, single-copy ACE-tRNA minicircles. The DNA bending protein used in this study was transcription factor A mitochondrial, TFAM, also known as ARS binding factor 2 protein, Abf2p (purification shown in Fig. S2). ACE-tRNA minicircles produced in this manner exhibit covalent closure as demonstrated by resistance to T5 exonuclease degradation, unless first linearized by an endonuclease (Fig. 1A and S3A).

Comparatively larger ACE-tRNA minicircles were produced using a commercially available minicircle parent plasmid and *E. coli* strain for production (System Biosciences and described in (38)). This system allows for production of minicircles in a manner similar to production of plasmid DNA with multiple copies of ACE-tRNA present per minicircle. Minicircles produced in this manner demonstrate T5 exonuclease resistance and DNA supercoiling (Fig. 1B and S3B).

While circular DNA is perhaps the conventional DNA structure envisioned for a DNA vector, linear DNA vectors with covalently closed ends have also shown the potential for utility as DNA therapeutics (35,39,40). Protelomerase from the N15 temperate phage (TelN) has been shown to recognize a 56 bp inverted repeat *telRL* DNA sequence and possess cleaving-joining activity. Functionally this means that a single *telRL* site is processed by TelN to separate DNA molecules containing *telR* and *telL* ends covalently closed by phosphodiester bonds (41,42). To produce closed end DNA threads (CEDT) minivectors using TelN, ACE-tRNA expression cassettes were cloned between *telRL* sites in a pUC57 Kan vector. TelN DNA substrate was produced by PCR with primers designed to flank the *telRL* sites such that the PCR product contains the *telRL* recognition sites for TelN. TelN was produced by recombinant expression and purified by Ni-NTA and CEX chromatography (Fig. S4). To decrease the TelN dependence of ACE-tRNA CEDT production, especially for the smaller 200 bp CEDTs, the ACE-tRNA expression cassettes were cloned back-to-back, sharing a common *telRL* site between the two cassettes (Fig. 1C). After generation via TelN reaction and further purification, as for the other minivectors, ACE-tRNA CEDTs exhibit exonuclease resistance (Fig. 1C and S3C).

### ACE-tRNA minivectors exhibit efficient PTC suppression in the HEK293T cells

To obtain a baseline understanding of the PTC suppression activity of the ACE-tRNA minivectors, both ACE-tRNA^Arg^_UGA_ and ACE-tRNA^Leu^_UGA_ expressing minivectors were tested, as the Arg CGA > UGA is the most common disease-causing PTC across all genetic diseases (43), and ACE-tRNA^Leu^_UGA_ was shown to efficiently suppress and recover channel function in excess of the therapeutic threshold for the four most common CF-causing *CFTR* PTCs (22). ACE-tRNA minivectors of several different sizes, ranging from 200 bp minivectors to 5000 bp plasmids, and consisting of several different DNA structures, including circular and linear, were co-transfected using Lipofectamine 2000 along with a previously described PTC reporter (16) into HEK293T cells (Fig. 2A and Fig. S5A-C) and the PTC suppression efficiency of each was determined by Dual-luciferase assay (Fig. 2B and S6). A scrambled ACE-tRNA cassette (S*) was used as a negative control, which should not be transcribed as it does not possess A-and B-boxes for type 2 Pol III transcription (44) as opposed to the active ACE-tRNA expression cassettes employed here. As we are ultimately interested in these minivectors as therapeutic vectors, we assessed their function on a per-mass basis as the implementation of most therapeutic DNA delivery modalities are determined by mass of DNA delivered, not moles of active DNA cargo.

The majority of ACE-tRNA minivectors tested displayed efficient PTC suppression following delivery, with the exception of the 200 bp minicircles and several annealed single-stranded minivectors, which did not display significantly more PTC suppression than the scrambled control vectors (Fig. 2B and S6). While both nicked and supercoiled 200 bp minicircle demonstrated significantly more efficient PTC suppression than the open circle minicircle (Fig. S7), they were still ∼15-fold less efficient at PTC suppression than the 400 bp minicircle. In contrast however, the 200 bp CEDTs did promote efficient PTC suppression, making these vectors, to our knowledge, the smallest functional expression vectors ever produced. To demonstrate that ACE-tRNA minivectors are competent to rescue PTCs in therapeutic target proteins, rescue of full-length protein expression for both Rep-1-R253X-Nluc (Fig. 2C and S8) and CFTR-W1282X-Nluc (Fig. 2D and S8) was assessed by in-gel-luminescence. The 850 bp 4x ACE-tRNA^Arg^_UGA_ and 4x ACE-tRNA^Leu^_UGA_ minivectors were co-transfected along with the Rep-1 or CFTR expression vectors in HEK293T cells. Total protein was resolved by SDS-PAGE and the Nluc tag luminescence was imaged with rescued protein amounts determined by band densitometry. Significant rescue of PTC-containing protein was apparent for both Rep-1 R253X (78 ± 3% of WT) and fully glycosylated CFTR W1282X C-band (84 ± 5% of WT).

### ACE-tRNAs encoded as minivectors display efficient delivery to human bronchial epithelial cells

To determine a summary of the impact of ACE-tRNA minivector DNA size and structure on PTC-suppression efficiency, we transfected (Lipofectamine 2000) and nucleofected (Lonza Nucleofector) the minivectors that had demonstrated function (Fig. 2B) into the well-characterized, immortalized, isogenic human bronchial epithelial cell line (16HBE14o-) containing a stably-expressing Stop-Go-Glow (SGG) PTC-suppression reporter (Fig. 3A and S5D-F). This PTC reporter cassette allows for high-throughput determination of PTC-suppression efficiency via a PTC-interrupted nanoluciferase (12,15,16), and cell-by-cell analysis (e.g. by flow cytometry) of PTC-suppression via a downstream mScarlet fluorescent reporter (Fig. S9). In all cases, the ACE-tRNA^Arg^_UGA_ and ACE-tRNA^Leu^_UGA_ minivectors displayed significantly higher PTC suppression efficiency by Nluc assay as compared to the respective scrambled controls (Fig. 3B). The 850 bp 4x ACE-tRNA minicircles when nucleofected set the upper limit for average PTC suppression efficiency with significantly (adjusted *p* value <0.0001) higher PTC suppression than all other conditions tested.

We have previously established an upper limit to the transfection efficiency of pDNA into 16HBE14o-cells as ∼40% (15). Here we show that the minivectors tested exhibit delivery efficiencies of up to ∼60%, as demonstrated by the mScarlet positive cell population (Fig. 3C and 3E). While minivectors demonstrate a larger mScarlet positive cell population as compared to pDNA, the median fluorescence intensities (MFIs) do not show a marked difference (Fig. S10). This indicates that while a greater percentage of the cell population is receiving minivector as compared to pDNA, the level of PTC suppression is not markedly higher.

### Optimized ACE-tRNA cassettes encoded as DNA minivectors display lower DNA-dependence for efficient PTC suppression

During preparation of this manuscript, a screening effort in the lab for multiple ACE-tRNA functional elements resulted in several optimized ACE-tRNA expression cassettes with improved PTC suppression efficiency (16). In principle, these improved expression cassettes should impart similar favorable characteristics when encoded as ACE-tRNA minivectors. To that end, the optimized (Opt) ACE-tRNA^Arg^_UGA_ expression cassette was encoded as a 200 bp CEDT (Fig. 4A) and the PTC suppression efficiency was compared with the original ACE-tRNA^Arg^_UGA_ at a series of DNA concentrations when co-transfected with the dual-luciferase PTC reporter (Fig. 2A). As expected, the OptACE-tRNA^Arg^_UGA_ resulted in ∼2-fold higher peak PTC suppression (Sup_max_ 1500 ± 200 a.u.) as compared to the original ACE-tRNA^Arg^_UGA_ (Sup_max_ 760 ± 125 a.u.) (Fig. 4B and S11) and ∼4-fold lower DNA delivery amount to reach the same normalized suppression ratio (630 a.u. = 18.75 ng Opt vs 75 ng original).

### Optimized ACE-tRNA cassettes encoded as DNA minivectors efficiently rescue the Rep-1 R253X PTC

The decreased DNA delivery dependence and higher peak suppression efficiency for the 200 bp CEDT OptACE-tRNA^Arg^_UGA_ carried over when assessing Rep-1 R253X PTC rescue. Rep-1 constructs (Fig. 4A) with C-terminal ALFA tag (45) were co-transfected into HEK293T cells along with variable amounts of ACE-tRNA minivector, and the amount of Rep-1 expressed was determined by western blot (Fig. 4C-D, S12). OptACE-tRNA^Arg^_UGA_ when delivered at a 4-fold lower amount rescued the same amount of Rep-1 R253X as compared to the original cassette, and when comparing the same amount of DNA delivered, both 850 bp 4x ACE-tRNA^Arg^_UGA_ and 200 bp OptACE-tRNA^Arg^_UGA_ rescued the same amount of Rep-1 R253X (85 ± 5%, Fig 4D), while 4x ACE-tRNA^Arg^_UGA_ pDNA displayed a diminished but still significant level of Rep-1 R253X rescue (45 ± 5%, Fig 4D), with all ACE-tRNA treated samples well above the therapeutic threshold for Rep-1 rescue of <10%, as CHM is predominately caused by a complete lack of the affected Rep-1 protein (46).

### ACE-tRNA cassettes encoded as DNA minivectors efficiently rescue mRNA and protein expression for an endogenously encoded CFTR W1282X PTC

The CFTR transcripts in p.W1282X-*CFTR* 16HBEge cells harboring this PTC in the genuine genomic locus (Fig. 5A) are sensitive to degradation through the NMD quality control pathway (47), which we and others have shown to be rescued by the action of ACE-tRNA-mediated PTC suppression (13–16,22). To determine the level of NMD inhibition exhibited by ACE-tRNA minivectors, we nucleofected ACE-tRNA^Leu^_UGA_ minivectors into W1282X-*CFTR* 16HBEge cells and assessed the level of *CFTR* mRNA by real-time qRT-PCR as described previously (15) (Fig. 5B). Both minivectors tested displayed significant rescue of W1282X-containing *CFTR* mRNA transcript as compared to the scrambled control with the 850 bp 4x ACE-tRNA^Leu^_UGA_ minicircle displaying WT-levels of *CFTR* mRNA (Fig. 5B). ACE-tRNA^Leu^_UGA_ encoding minivectors were chosen to rescue the W1282X *CFTR* variant as this protein variant has been shown to provide near-WT CFTR channel function (15,48) and ACE-tRNA^Leu^_UGA_ has demonstrated functional rescue in excess of the therapeutic threshold for 4 of the most common PTC-associated *CFTR* variants harbored by people with CF (pwCF) (22). When assessed by western blot the 4x 850 bp ACE-tRNA^Leu^_UGA_ minicircle rescued near-WT levels of CFTR C-band (83 ± 6%), while the 200 bp ACE-tRNA^Leu^_UGA_ CEDT displayed significant rescue of CFTR C-band (20 ± 6%), which while relatively modest, in combination with CFTR modulators (22) is well above the therapeutic threshold for CFTR rescue (Fig. 5C-D, S13).

**Figure 5.**
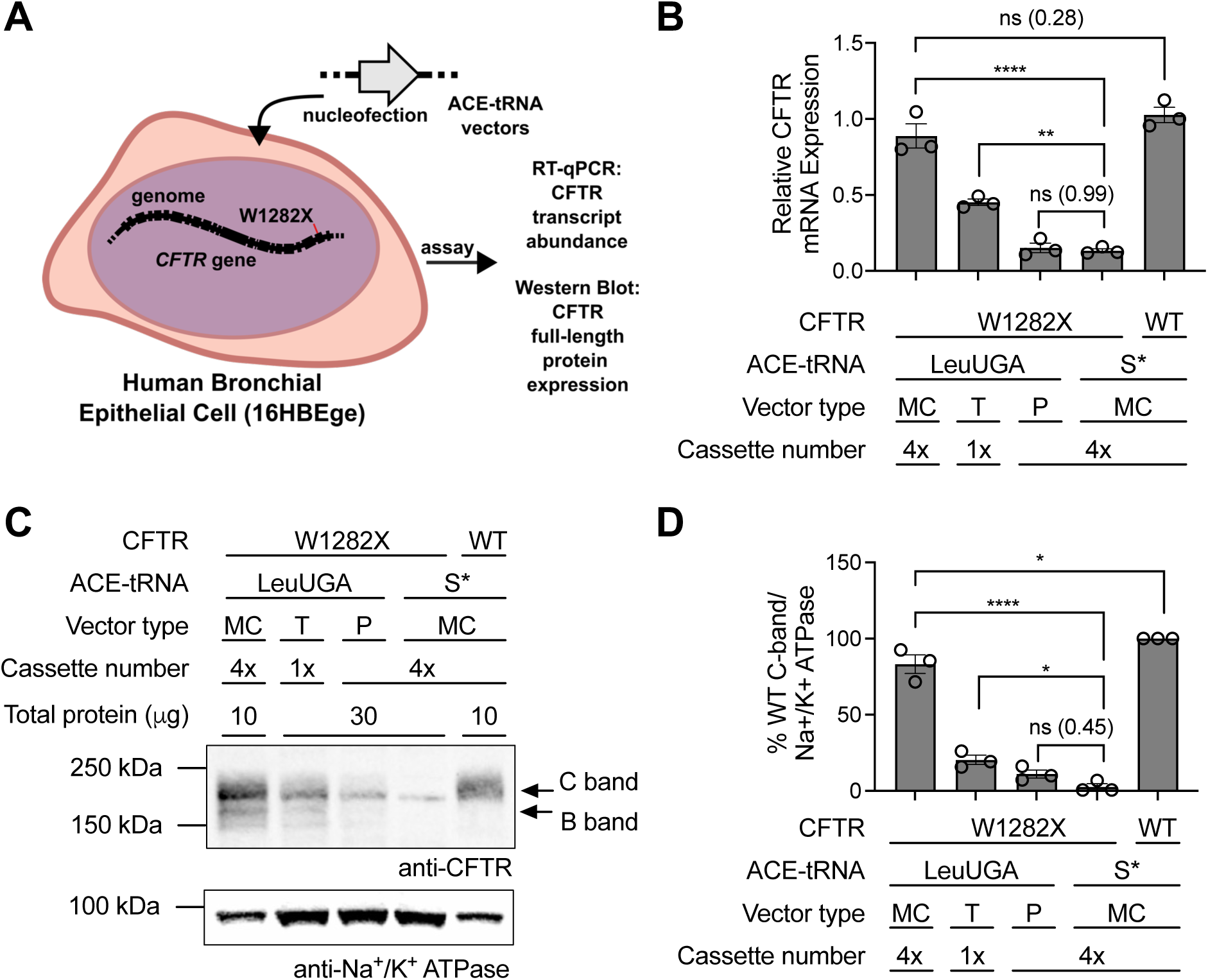
ACE-tRNAs encoded as DNA minivectors can rescue expression of the genomically encoded *CFTR* p.W1282X variant in 16HBEge cells. (**A**) Diagram of experimental setup with ACE-tRNA vectors delivered by nucleofection into 16HBEge cells containing the *CFTR* p.W1282X variant. (**B**) RT-qPCR analysis of relative CFTR expression in 16HBEge cells nucleofected (n = 3) with the indicated ACE-tRNA vector. (**C**) Representative western blot image showing *CFTR* p.W1282X rescue following co-transfection with the indicated ACE-tRNA vector. WT indicates that the parental 16HBE14o-cell line was used. (**D**) Quantified band densitometry data for the CFTR western blots (n = 3) with data shown as the percent of WT C-band (fully glycosylated form of CFTR) intensity normalized to the Na+/K+-ATPase loading control. Abbreviations: Minicircles, MC; Closed End DNA Threads, T; Plasmid DNA, P; Scrambled ACE-tRNA Control, S*. Data are presented as the mean ± SEM with significance determined by one-way ANOVA and Tukey’s post-hoc test, where **** *p* < 0.0001, and ns denotes not significantly different (*p* value shown for each ns and *).

**Figure 6.**
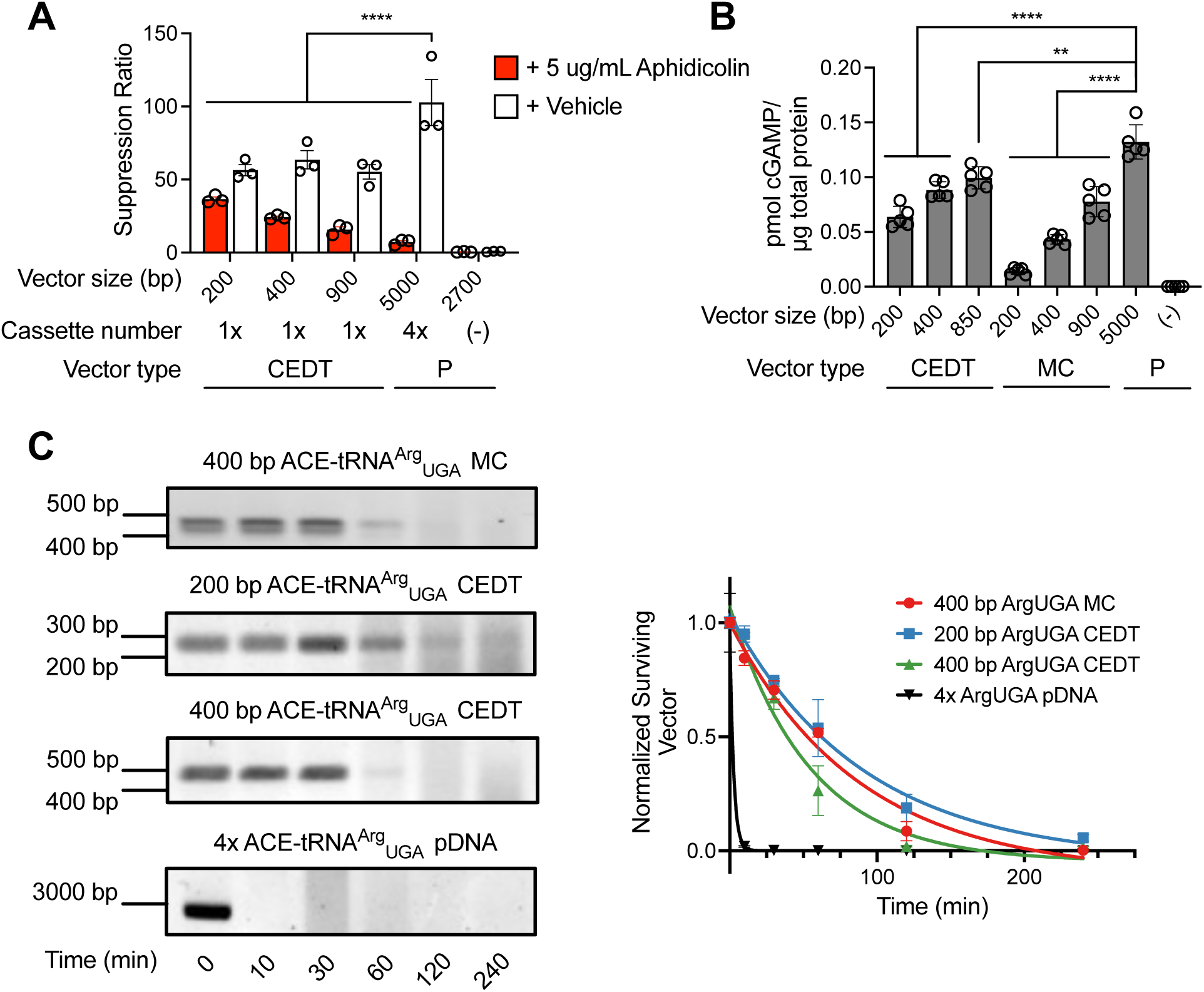
ACE-tRNA DNA minivectors exhibit better bioavailability, lower innate immune response induction, and better biostability as compared to pDNA. (**A**) 16HBE14o-cells containing the genomically encoded SGG PTC reporter were cultured and transfected either in the presence of the antimitotic agent aphidicolin or the DMSO vehicle with ACE-tRNA CEDT or pDNA (n = 3). NLuc-PTC suppression was used to demonstrate levels of PTC rescue for the various vectors (**B**) HeLa cells were transfected with ACE-tRNA MC, CEDT, or pDNA and after 6 hours in culture, were lysed, and assayed for cyclic GMP-AMP (cGAMP) levels by competitive ELISA (n = 5). The level of cGAMP present demonstrates activation of the cGAS-STING innate immune response pathway. (**C**) ACE-tRNAs encoded as either MC, CEDT, or pDNA (P) were subjected to incubation in human blood serum at 37 °C for the times indicated. After treatment vector DNA was purified by phenol/chloroform extraction, precipitated, and the amount of residual intact DNA was assessed by agarose gel electrophoresis and band densitometry (n = 3). Band densitometry data was fit to a one phase decay using non-linear regression, which is shown on the graph. Data are presented as the mean ± SEM with significance determined by one-way ANOVA and Tukey’s post-hoc test, where ** ** *p* < 0.01 and **** *p* < 0.0001.

### ACE-tRNA DNA minivectors display favorable bioavailability and biostability characteristics as compared to pDNA

Actively dividing cell lines like the ones employed in this study are mitotically active with each cell division resulting in the breakdown and re-formation of the nuclear membrane, allowing for DNA vectors to reach the nucleoplasm (49,50). In contrast, many of the targeted cell types for PTC-associated disorders are post-mitotic, meaning the DNA cargo must be able to reach the nucleoplasm independent of nuclear breakdown. To test the function of ACE-tRNA minivectors in a system similar to a post-mitotic cell, we treated 16HBE14o-cells containing the SGG transgene with the antimitotic drug aphidicolin, which has been shown to arrest mitosis without affecting transcription or translation (51). While pre-treatment with aphidicolin decreased the PTC suppression efficiency of pDNA by ∼14-fold, the efficiency of 200 bp CEDT DNA was only decreased by ∼1.5-fold, with 400 bp and 900 bp CEDT displaying intermediate effects, with efficiency being decreased by ∼2.5 and ∼3.5 respectively (Fig. 6A). The minimal impact of cell cycle arrest on the 200 bp CEDT ACE-tRNA function may reflect previous findings that DNA fragments as small as 200 bp can freely diffuse in the cytoplasm (52) and enter nuclear pores (50,53), meaning that nuclear breakdown would not be expected to be necessary for entry into the nucleoplasm.

DNA vectors face further challenges as therapeutic cargo, as the cyclic GMP-AMP synthase – stimulator of interferon genes (cGAS-STING) innate immune response pathway can respond to introduction of cytoplasmic double-stranded DNA (dsDNA) resulting in expression of inflammatory genes (54,55). The cytosolic cGAS enzyme has evolved to detect >45 bp long (in humans) stretches of cytosolic DNA and catalyze production of the second messenger 2’-5’/3’-5’ cyclic-GMP-AMP (cGAMP), which activates the receptor STING, ultimately leading to activation of type I interferon signaling (56). While this system is targeted to sense pathogen replication and abnormal cytosolic self-DNA, it would also be predicted to be activated in response to intracellular delivery of therapeutic vector DNA. To test the impact of vector DNA size on cGAS activation, we measured production of cGAMP in response to transfection of DNA vectors into HeLa cells, which have previously been demonstrated to possess a robust cGAS response (57). All minivectors tested here demonstrated a significantly lower level of cGAMP production than pDNA (Fig. 6B), with 200 bp CEDT and 200 bp minicircle each displaying a ∼2-fold lower and ∼9-fold lower cGAMP production than pDNA respectively. The discrepancy in cGAS activation despite similar polymer sizes between the two vectors may be related to DNA structure as these minivectors also display discrepancies in PTC suppression efficiency, which is presumed to be related to different efficiencies of transcription of the ACE-tRNA cargo from each of these vectors (Fig. 2B). It should also be noted that even when employing pDNA as a therapeutic cargo, cGAS-STING activation has not limited the application of DNA vectors for clinical development (58,59). Given the significantly lower activation of cGAS by minivector DNA as compared to pDNA, it would be expected that the innate immune response to ACE-tRNA therapeutic minivectors would not be cause for safety concerns.

In the absence of polycation carriers, therapeutic vector DNA must overcome enzymatic degradation in the bloodstream for effective delivery to affected tissue in the body (27). To that end, we tested the stability of ACE-tRNA minivector and plasmid DNA in human serum.

Samples of DNA vectors were incubated at 37 °C in 100% human serum, at time points indicated (Fig. 6C and S13) samples were removed and the DNA purified by phenol/chloroform extraction. Residual DNA was resolved by agarose gel electrophoresis and the amount of vector remaining was determined by band densitometry. While ACE-tRNA minivector DNA displayed a half-life of 45-60 minutes in the assay, no residual supercoiled ACE-tRNA pDNA remained after 10 minutes of treatment, with the supercoiled pDNA displaying a half-life of ∼2 minutes (Fig. 6C). The longer lifetime of minivector DNA in human serum has been shown previously (60), which indicates a higher likelihood of successful delivery to affected tissue *in vivo* without encapsulating carriers.

Taken together, these findings indicate that ACE-tRNA minivectors possess favorable characteristics as therapeutic DNA vectors as compared to conventional plasmid DNA vectors.

## DISCUSSION

The burgeoning field of sup-tRNA therapeutics has been led by the development of RNA and AAV viral-vector based delivery approaches (13,14). Due to potential therapeutic hurdles related to frequent redosing of RNA therapeutics and immune responses limiting the use of AAV viral vectors, we wanted to broaden the portfolio of ACE-tRNA therapeutic vectors to include minimal, non-viral, DNA vectors. Functional ACE-tRNA expression cassettes can be composed of DNA sequences as short as 125 bp (8). DNA vectors this small possessing transcriptional activity have not been investigated previously and as such we wanted to probe the limits of DNA size and shape for functional expression of ACE-tRNAs, as these minimal DNA vectors could potentially possess improved attributes related to therapeutic delivery or function.

Non-viral DNA vectors must overcome several extracellular and intracellular barriers before efficient expression of their therapeutic cargo can be carried out (27,49,61–63). The DNA vector must survive the extracellular environment long enough for cellular association and uptake to take place (64,65), it must then undergo endosomal escape (66,67) in a spatiotemporially controlled manner (68), undergo proper intracellular trafficking (69–74) and dissociate from any polycation carrier, undergo nuclear import (75–77), and incorporate into active chromatin for sustained transcription (78,79). Of these steps, diffusion in the cytosol and nuclear import are believed to be rate-limiting determinants of transgene delivery (80,81). Here we demonstrate that ACE-tRNA minivectors are able to effectively transit all of these steps when delivered either by the “gold standard” of lipoplexes (Lipofectamine 2000) (82) or by an orthogonal gene delivery approach, electroporation. Even in mitotically arrested cells which do not go through nuclear membrane breakdown, 200 bp CEDT ACE-tRNAs are able to effectively transit the cell and enter the nucleus (Fig. 6A). Given the somewhat unique physical characteristics of ACE-tRNA minivectors, existing delivery modalities may need to be heavily adapted for optimal delivery of ACE-tRNA minivectors as has been previously demonstrated for delivery of other small nucleotide cargos (49). Nonetheless, given that both lipofection and electroporation worked for ACE-tRNA minivector delivery, it is highly likely that the wide range of previously developed DNA delivery modalities can be adapted for use with ACE-tRNA minivectors.

Strikingly, while the 200 bp CEDT supported expression of a functional ACE-tRNA, the 200 bp minicircle did not. These results likely reflect differences in DNA structure, with the 200 bp circle experiencing steric strain (83), which the linear vector does not experience. Structural characterization of the RNA pol III transcriptional machinery assembled on target DNA demonstrates a bend in the modeled DNA backbone (Fig S1C) (84), which may be incompatible with effective transcription from the 200 bp ACE-tRNA minicircle. This could also potentially be due to less room to distribute topological tension resulting from unwinding to enable transcription (85), or while unlikely, could be related to differences in cell entry or trafficking.

Previous work has shown that linear minivectors (also referred to as MIDGE vectors) generally demonstrate higher levels of transgene expression (35,86). Here we report similar or slightly lower levels of ACE-tRNA expression from linear vectors (CEDTs), even at higher molar quantities of transgene delivered. This discrepancy could reflect differences in transcription factor scanning or the efficiency of transcription initiation or termination (87,88) on the particularly small DNA vectors employed here. The efficiency of vector delivery by electroporation has been shown to be independent of vector length for vectors between ∼400-4500 bp, however smaller vectors did display lower function per mole (85), which is consistent with our findings here.

While the 850 bp 4x ACE-tRNA minicircles demonstrated the highest nonsense suppression efficiency of all vectors tested (Fig. 2 and 3), which may reflect the higher transfection efficiency of supercoiled DNA vectors (40,89), or the influence of supercoiling on transcription factor binding and transcription initiation in the promoter region (90). Ultimately this vector format is unlikely to see success as a therapeutic due to poor production yields (∼30 μg pure minicircle per L of *E. coli* culture), a laborious purification scheme to remove parent plasmid, and additional cGMP challenges due to the presence of endotoxin (89). The minicircle production system does not appear to be particularly amenable to production of ACE-tRNA minicircles, perhaps due to the small size of the therapeutic cargo. The small size of the product minivector would be expected to place constraints on the recombinase system not previously encountered in the production of larger RNA pol II expression cassette-containing minicircles and may explain the poor minicircle production yields noted for ACE-tRNA minicircles. Both *in vitro* methods (Fig. 1A and 1C) resulted in higher yield production of minivectors with yields on the order of 1 mg of minivector product per 100 mL of PCR product produced. While the methods employed here were sufficient to produce adequate amounts of minivector for *in vitro* investigations in PTC-containing cell models, cGMP production will likely require establishing a more streamlined process for large scale production of therapeutic ACE-tRNA minivectors.

DNA minivectors have been demonstrated to possess a number of attractive qualities *in vivo*, and while ACE-tRNA minivectors have not yet been tested in this manner, it would be anticipated that they will share many of the same qualities. Previous studies have demonstrated enhanced transgene expression from minicircle as compared to pDNA in liver (91–93), heart (94,95), skeletal muscle (95), and lung (96). While pDNA initially supports expression of high levels of transgene product, this effect quickly diminishes over a period of days (91,97–101).

This effect appears to be a result of silencing of transcription from the pDNA vectors (102–104). In contrast, minivector DNA supports long term expression of the target transgene for periods of up to 56 days in mouse lung (102), and >115 days in mouse liver (91). As minivector administration is typically less immunogenic than delivery of viral vectors (105,106), repeat administration is allowed, a critical feature as gene therapies for lung diseases will likely required repeat dosing several times per year for the lifetime of the patient (96). Beyond the improved nuclease resistance of the ACE-tRNA minivectors demonstrated here (Fig. 6C), minivectors have demonstrated better mechanical stability allowing for increased delivery of intact vector by physical methods such as nebulization for delivery to the lungs (107,108).

Beyond the minivectors themselves, the ACE-tRNA cargo has demonstrated considerable advantages as a gene therapy modality. While for any monogenetic disorder, total gene replacement represents a mutation agnostic approach, every disorder will require development of an independent therapeutic cargo. In contrast to total gene replacement, ACE-tRNAs are gene agnostic, with the only dependence being upon the PTC identity (UAA, UAG, or UGA) and the amino acid encoded. We have further demonstrated the potential of an amino acid agnostic ACE-tRNA platform approach providing therapeutic coverage for a broad swath of PTC sites and genetic diseases like CF (22). If conditions for effective delivery of ACE-tRNA minivectors to affected lungs for the treatment of CF can be ascertained, it is likely that PTC-associated disorders in similar cell types like primary ciliary dyskinesia will be amenable to treatment using very similar or identical methods. Further, ACE-tRNAs maintain native regulation of the PTC-containing gene (15), important for diseases like CDKL5 deficiency disorder (18), for which overexpression of the target protein can lead to deleterious effects.

While natural termination codon (NTC) readthrough has been a reasonable concern when employing ACE-tRNAs as a therapeutic approach, all available data indicate that while ACE-tRNAs display efficacious readthrough at PTCs, they are considerably less efficient at NTC readthrough (12–14). While the mechanisms involved are complicated, these findings likely reflect that NTCs simply have evolved for efficient translation termination, while PTCs have not. The safeguards in place to deal with spurious NTC readthrough include in-frame downstream stop codons (109,110), the stop codon nucleotide influence on readthrough (111,112), the distance to the polyA binding protein (113,114), polyLys tracts leading to ubiquitination and degradation (115), non-stop decay (116), and readthrough of only a single stop codon (UAA, UAG, or UGA) by ACE-tRNAs (12). With these cellular safeguards in place, it has become clear that a therapeutic window should exist for delivery of ACE-tRNAs consistent with both the efficacy and the safety of this approach.

While non-viral DNA vector delivery approaches have demonstrated lower levels of transduction in airways, lack cell type specific targeting, and have shown an inability to transfect nondividing cells when compared with viral vectors (117), ACE-tRNA minivectors display favorable characteristics when compared to pDNA. Minivectors have demonstrated improved transfer efficiency when delivered by polyplex (118) or electroporation (85,119), and while the delivery methods employed here were optimized for pDNA delivery, future improvements could include delivery modalities tailored to the ACE-tRNA minivectors themselves (49,120). Further, one could envision altering the *in vitro* production methods to incorporate cellular or intracellular targeting moieties into the ACE-tRNA minivector to improve delivery efficiency. Receptor-mediated targeting, nuclear targeting, and improved endosomal escape could all be targeted to bridge the gap between the efficacy of viral and non-viral approaches for ACE-tRNA delivery *in vivo*.

Here we report that ACE-tRNA minivectors as small as 200 bp can efficiently suppress PTCs in several genes related to PTC-associated monogenetic diseases. Further, we demonstrate that ACE-tRNA minivectors display improved transfer efficiency, better bioavailability, a lower innate immune burden, and better biostability as compared to conventional pDNA vectors. With these favorable characteristics, ACE-tRNA minivectors broaden the therapeutic approaches available for delivery of sup-tRNA delivery for treating human diseases.

## ACKNOWLEDGMENTS

We thank members of the Lueck Laboratory for reading and editing the manuscript and constructive discussion throughout the study. We would like to thank the Cystic Fibrosis Foundation Therapeutics Lab and Dr. Hillary Valley for providing the 16HBE14ge cell lines used in this study. This work was supported by Cystic Fibrosis Foundation Postdoctoral Fellowship (PORTER20F0) and The Throssell and Hillier Families, and Choroideremia Research Foundation Research Award to J.J.P., a NIH grant (GM151450) to D.A.D. and an NIH grant (HL153988) and Vertex Innovation Cystic Fibrosis Research Award to J.D.L.

## AUTHOR CONTRIBUTIONS

J.J.P., W.K., D.A.D., and J.D.L. designed the study. J.J.P., W.K., E.G.S., Z.C., T.C., J.G., V.S., J.H., J.L., and L.P. and performed experiments. J.J.P., E.G.S., T.C., and W.K. analyzed the data and constructed the figures. J.J.P. and J.D.L. wrote the manuscript. All authors read and revised the manuscript.

## DATA AVAILIBILITY

The data underlying this article are available in the article and in its online supplementary material.

