## Supplemental Information for "Anticodon Edited Transfer RNAs (ACE-tRNAs) Encoded as Therapeutic Nonviral Minimal DNA Vectors"

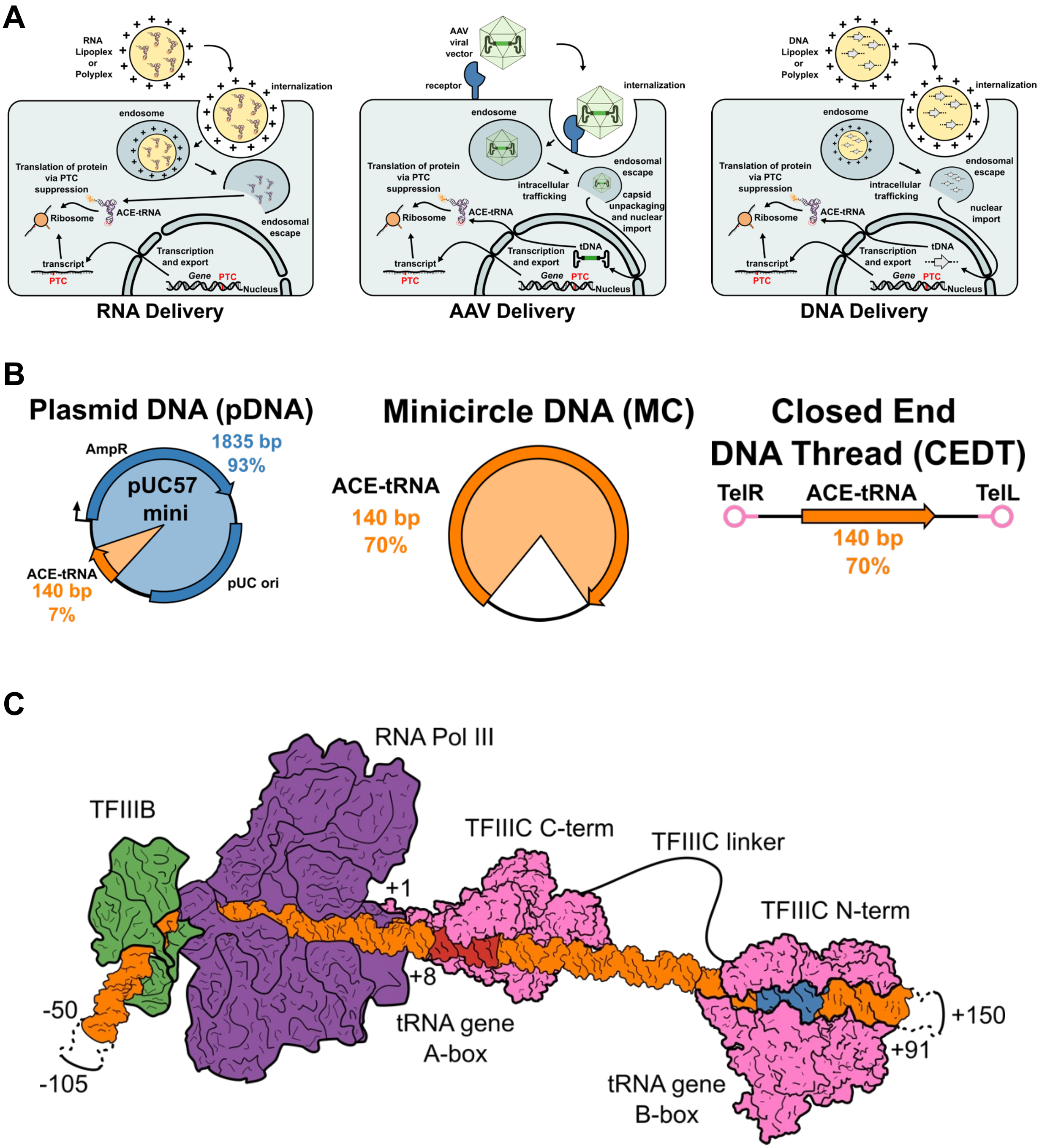


**Supplemental Figure 1. ACE-tRNA vectors and assembly of RNA Pol III transcription machinery on an ACE-tRNA minivector.** (**A**) Several methods have been used for ACE-tRNA delivery to cells/tissues for treatment of PTC-affected genes. ACE-tRNAs can be delivered to cells as lipoplex or polyplex formulations, which are internalized into endosomes, following endosomal escape the ACE-tRNAs are charged by their endogenous aminoacyl-tRNA synthetase (aaRS), and function to suppress PTCs on the ribosome, rescuing expression of PTC-containing proteins from mRNA transcripts expressed and exported from the nucleus. ACE-tRNAs can also be encoded in AAV viral vectors, which have demonstrated safety and efficacy as gene therapy vectors. AAVs have been engineered with certain tropisms through targeting of receptors for internalization, possess effective endosomal escape and nuclear import capabilities, and allow for expression of therapeutic cargo (in this case the tDNA) on a timescale of months to years. ACE-tRNAs transcribed in the nucleus from the tDNA are exported and processed to the mature ACE-tRNA form, charged by the aaRS and rescue expression of PTC-containing proteins. ACE-tRNAs can also be delivered as DNA vectors via lipoplex or polyplex formulations, following much the same path as delivery of the AAV viral vector and expression of the ACE-tRNA cargo. Internalization, endosomal escape, and nuclear trafficking are points of potential optimization to increase delivery of DNA vectors. Expression of ACE-tRNAs from the tDNA is expected to persist for the lifetime of the cell for continual PTC suppression activity. (**B**) Anatomy of plasmid DNA vectors containing both the plasmid DNA backbone (AmpR antibiotic resistance marker, and pUC origin of replication) and the therapeutic cargo DNA (ACE-tRNA). Even for the small pDNA vector pUC57 mini containing a single copy of the ACE-tRNA, the plasmid backbone DNA makes up much of the DNA vector (93%) while the ACE-tRNA active cargo is only a small fraction (7%). When the ACE-tRNA is encoded in a minivector, whether minicircle (MC) or closed end DNA thread (CEDT), the ACE-tRNA cargo makes up a much larger percentage (70%). (**C**) ACE-tRNAs expressed from DNA vectors (tDNA) must be effectively transcribed by RNA pol III. A stretch of CEDT minivector DNA containing an ACE-tRNA is modeled here along with assembled RNA Pol III machinery composed of transcription factor IIIC (TFIIIC), transcription factor III B (TFIIIB), and RNA Pol III. The tRNA A-box and B-box represent ACE-tRNA intragenic sequences which serve as binding sites for TFIIIC, which initiates assembly of the transcription complex. The model shown here is derived from structures presented in (1,2).

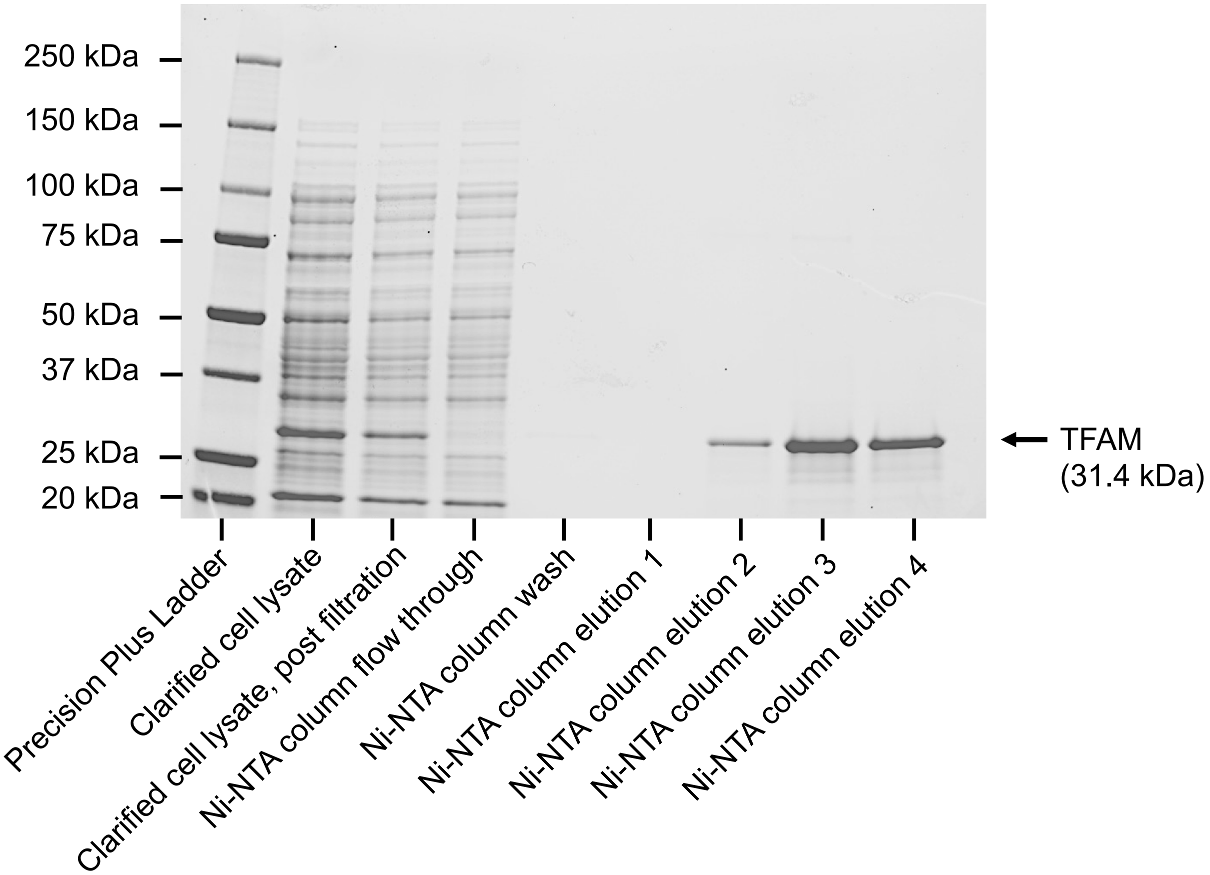


**Supplementary Figure 2. Expression and purification of transcription factor A mitochondrial (TFAM) for production of minicircle DNA.** The TFAM cDNA sequence containing an N-terminal 6xHis tag for expression was cloned in a pET28a(+) vector (sequence below). The TFAM pET28a(+) expression vector was transformed into NiCo21(DE3) competent *E. coli* and expressed in ZYM-505 media (3) and the cells were lysed, clarified, and the clarified lysate was subjected to Ni-NTA purification as per the methods section. Following purification 10 μL aliquots of the above samples were resolved via 4-12% SDS-PAGE electrophoresis and stained with coomassie blue.

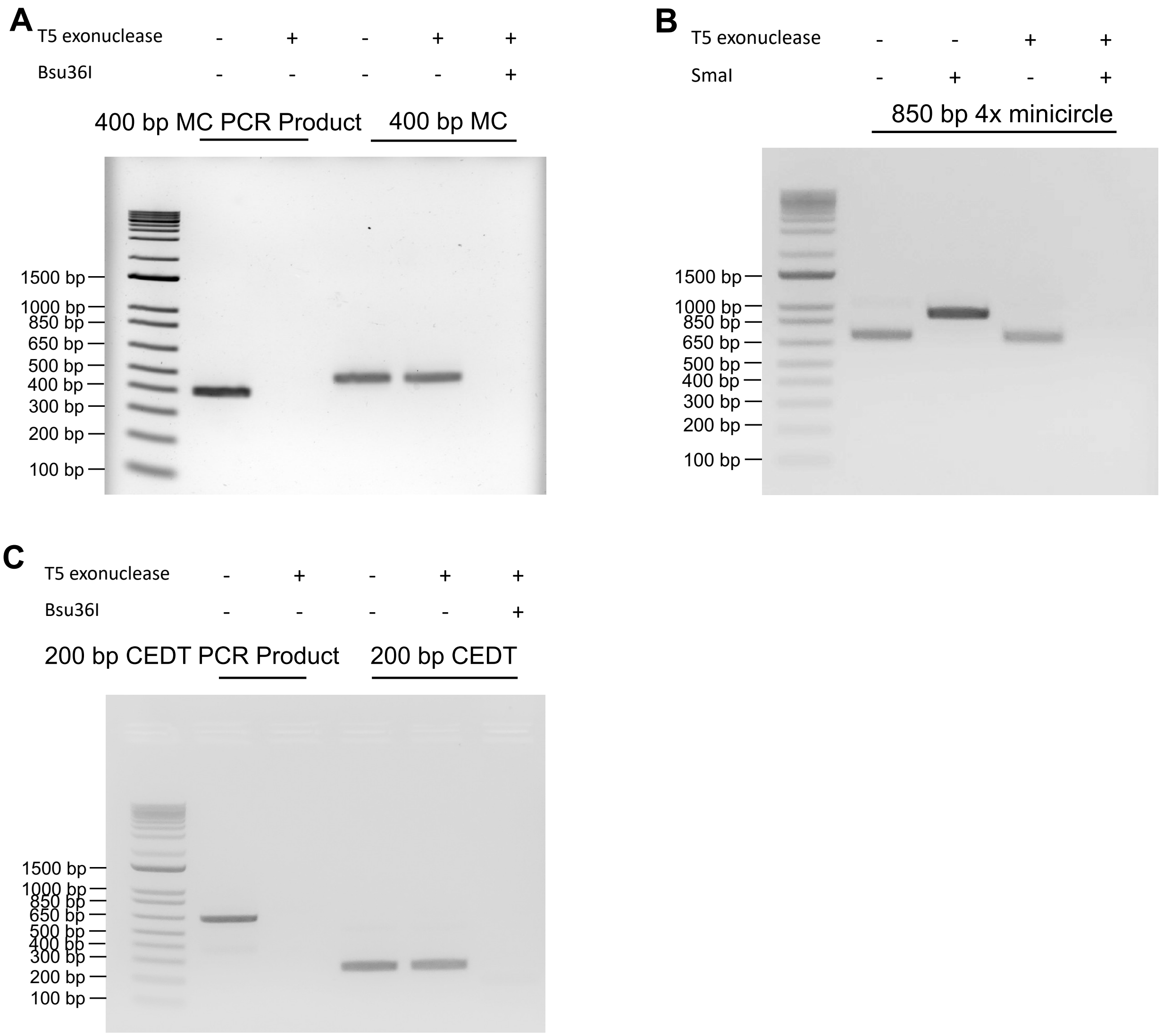


**Supplemental Figure 3.** Full agarose gel images of DNA minivector gels from Figure 1.

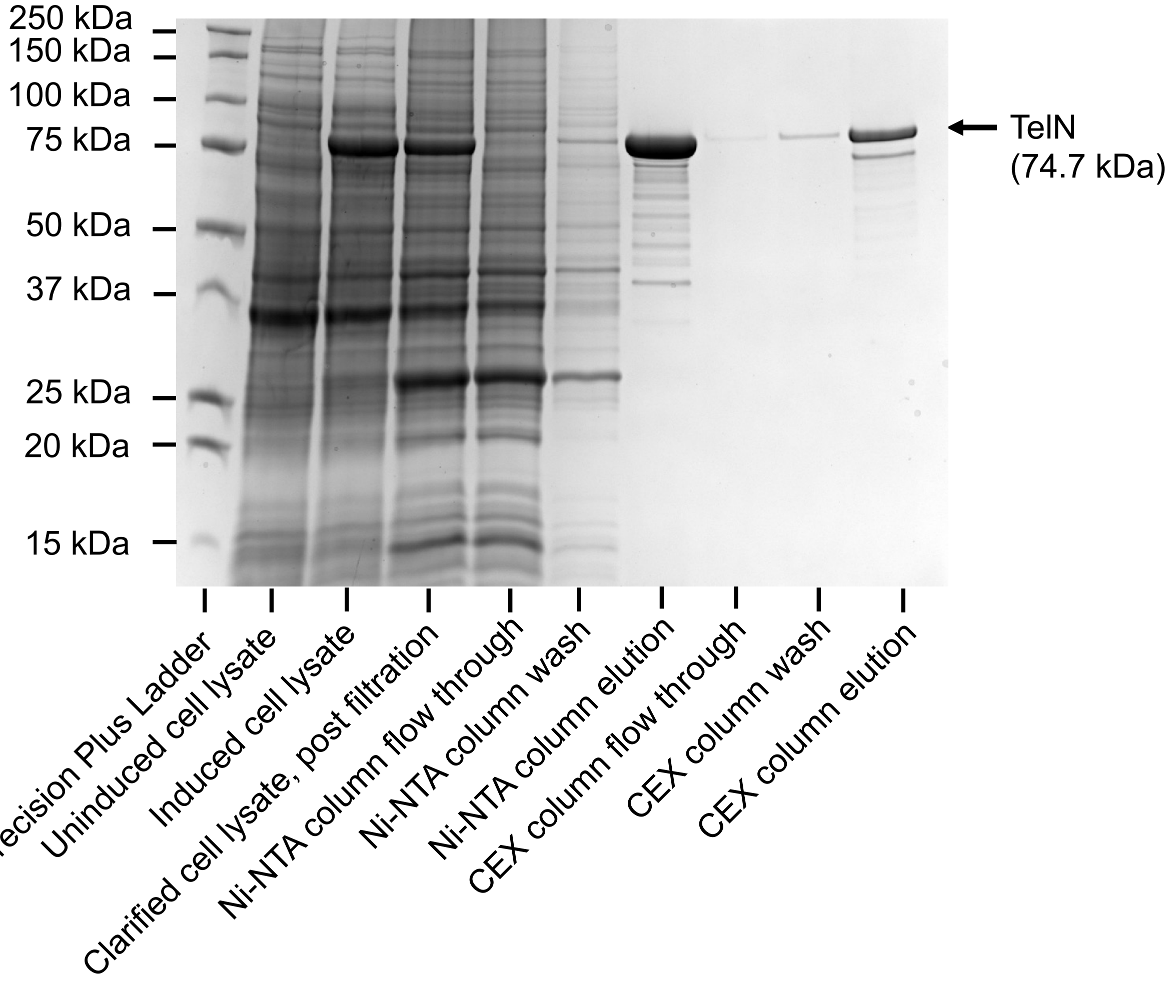


**Supplementary Figure 4. Expression and purification of protelomerase from *E. coli* temperate phage N15 (TelN) for production of CEDT minivector DNA.** The TelN CDS sequence containing an N-terminal 6xHis tag for expression was cloned in a pET21a(+) vector (sequence below). The TelN pET21a(+) expression vector was transformed into NiCo21(DE3) competent *E. coli* and expressed in ZYM-505 media (3) and the cells were lysed, clarified, and the clarified lysate was subjected to Ni-NTA purification as per the methods section. Following elution in low salt Ni-NTA elution buffer, the eluted protein was subjected to purification on a sulfopropyl-agarose cation exchange resin column. Following purification 10 µL aliquots of the above samples were resolved via 4-20% SDS-PAGE electrophoresis and stained with coomassie blue.

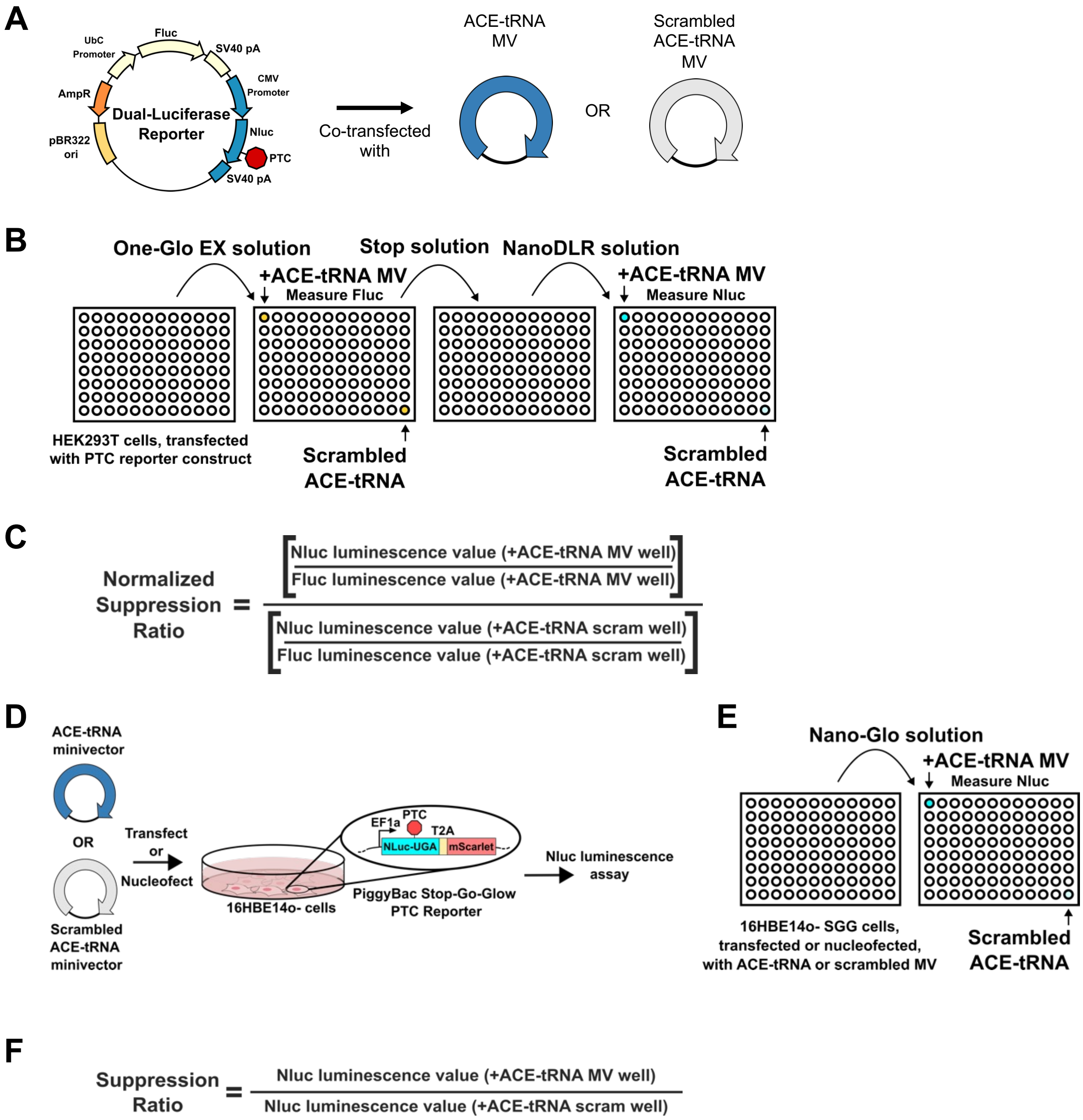


**Supplemental Figure 5. PTC reporter constructs and PTC suppression assay scheme** (**A**) The pNanoRePorter 2.0 PTC reporter plasmid (4) contains a firefly luciferase (Fluc) expression cassette for transfection normalization and a PTC-interrupted nanoluciferase (Nluc) expression cassette for high-throughput readout of PTC suppression. Minivector constructs containing scrambled-ACE-tRNA sequences are used as a negative control as they contain no active ACE-tRNA and thus only report on basal cellular PTC suppression under the conditions used for minivector delivery. (**B**) Following co-transfection of the PTC reporter plasmid and a scrambled-ACE-tRNA (lower right well) and the PTC reporter and an ACE-tRNA-containing minivector (upper left well) into HEK293T cells in 96-well format, the plates were maintained in a $CO_2$ cell culture incubator for 24 hours. After incubation, the media was aspirated, ONE-Glo Ex solution

(Nano-Glo Dual-Luciferase Assay Reporter Kit, Promega) containing Fluc substrate was added to the wells, and Fluc luminescence of each well was assayed by plate reader. After the Fluc luminescence was assayed, Stop solution was added to each well to quench Fluc luminescence. After Fluc luminescence was quenched, NanoDLR solution was added to each well and the Nluc luminescence was assayed by plate reader. High Nluc luminescence is seen for wells co-transfected with reporter constructs and minivectors containing ACE-tRNAs, while low Nluc luminescence is seen for wells co-transfected with reporter constructs and minivectors containing scrambled-ACE-tRNA. (**C**) The normalized suppression ratio for each minivector is computed based on the equation shown here. Every plate contains several wells co-transfected with the PTC reporter and a scrambled-ACE-tRNA control to normalize PTC suppression for all wells in that plate to account for basal cellular PTC suppression. (**D**) ACE-tRNA minivectors or scrambled-ACE-tRNA minivectors were transfected or nucleofected into 16HBE14o- cells stably expressing a Stop-Go-Glow (SGG) construct containing a PTC-interrupted Nluc expression cassette. As above, minivector constructs containing scrambled-ACE-tRNA sequences are used as a negative control as they contain no active ACE-tRNA and thus only report on basal cellular PTC suppression under the conditions used for minivector delivery. (**E**) Similar to as outlined above, the SGG cell lines were subjected to the Nano-Glo reagent (Nano-Glo Luciferase Assay System, Promega) to determine the PTC-suppression for either the ACE-tRNA MV treated or scrambled-ACE-tRNA treated wells and the luminescence was measured via plate reader. High Nluc luminescence is seen for wells treated with minivectors containing ACE-tRNAs, while low Nluc luminescence is seen for wells treated with minivectors containing scrambled-ACE-tRNA. (**F**) The suppression ratio for each minivector is computed based on the equation shown here. Every plate contains several wells treated with a scrambled-ACE-tRNA control to normalize PTC suppression for all wells in that plate to account for basal cellular PTC suppression.

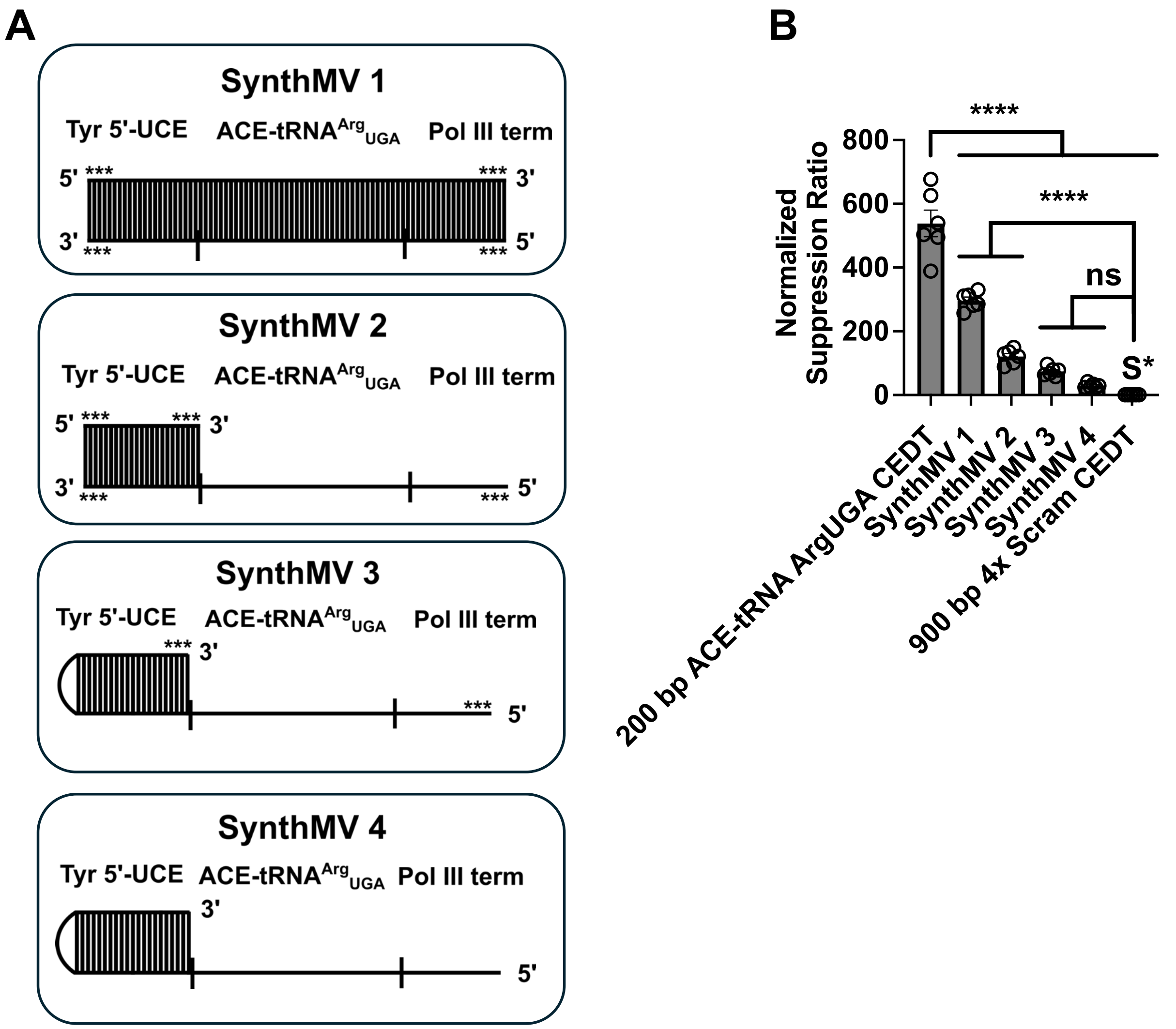


**Supplemental Figure 6. ACE-tRNA minivectors produced by solid-phase oligonucleotide synthesis demonstrate PTC suppression.** (**A**) A series of single stranded annealed ACE-tRNA minivectors were produced by solid-phase oligonucleotide synthesis (Integrated DNA Technologies). The components of the ACE-tRNA$^{Arg}_{UGA}$ expression cassette include the 5'-upstream control element (5'-UCE) derived from the 55 bp upstream of the human tRNA-Tyr-GTA-5-1, the ACE-tRNA$^{Arg}_{UGA}$ tRNA sequence, and the RNA Pol III transcription terminator sequence. Vertical lines represent base-paired regions of the SynthMV, 5' and 3' DNA ends are denoted, *** denotes that the last three nucleotides on that end of the DNA are linked by exonuclease-resistant phosphorothioate bonds. SynthMV 1 and 2 are composed of two independent complimentary oligos which are annealed to each other, while SynthMV 3 and 4 are composed of a single annealed self-complimentary oligo. (**B**) After annealing the complimentary SynthMV elements, the SynthMV were co-transfected along with a plasmid containing a PTC-interrupted nanoluciferase (Nluc-UGA) expression cassette and a firefly luciferase (Fluc) expression cassette, which serves as a transfection control (Fig 1A) using lipofectamine 2000 into the HEK293T cell line. The graph displays results of dual luciferase assays (n = 6) for each minivector. The normalized suppression ratio shown here is calculated from the equation (PTC-NanoLuciferase luminescence [+ACE-tRNA]/Firefly luminescence)/(PTC-Nanoluciferase luminescence [no ACE-tRNA]/Firefly luminescence). S*

denotes a minivector containing a scrambled ACE-tRNA, which serves as a negative control. Data are presented as the mean ± SEM with significance determined by one-way ANOVA and Tukey’s post-hoc test, where **** $p < 0.0001$, and ns denotes not significantly different.

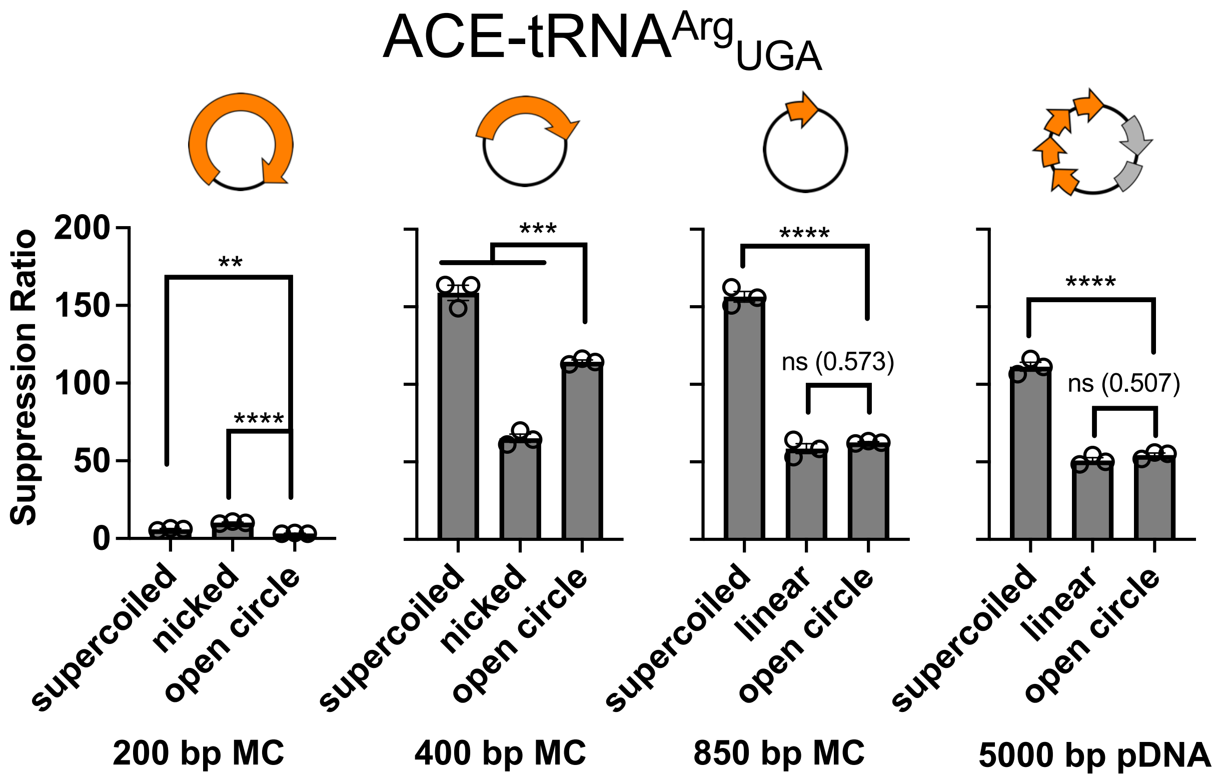


**Supplemental Figure 7. Supercoiled ACE-tRNA vectors support more efficient PTC suppression than open circle vectors.** Supercoiled 200 bp and 400 bp ACE-tRNA$^{Arg}_{UGA}$ minicircles were produced from open circle minicircles using the methods outlined in (5) and the nicking endonuclease Nt.AlwI. Minicircles denoted 'nicked' contained a single strand nick, while vectors denoted 'linear' were linearized by restriction endonuclease digestion (XmaI). Both were dephosphorylated with shrimp alkaline phosphatase after nuclease processing, and all vectors presented here were purified in triplicate by agarose gel extraction. The vectors as shown above were transfected into 16HBE14o- cells containing the stably integrated SGG v3 reporter (Fig. 3A) using Lipofectamine 2000. At a timepoint 24 hours post-transfection the cell media was removed and the Nluc signal for each well was determined by plate reader. The suppression ratio shown here is calculated from the equation (PTC-NanoLuciferase luminescence [+ACE-tRNA])/(PTC-Nanoluciferase luminescence [no ACE-tRNA]). Data are presented as the mean ± SEM with significance determined by one-way ANOVA and Tukey's post-hoc test, where ** $p < 0.01$, *** $p < 0.001$, **** $p < 0.0001$, and ns denotes not significantly different with $p$ values shown in this case.

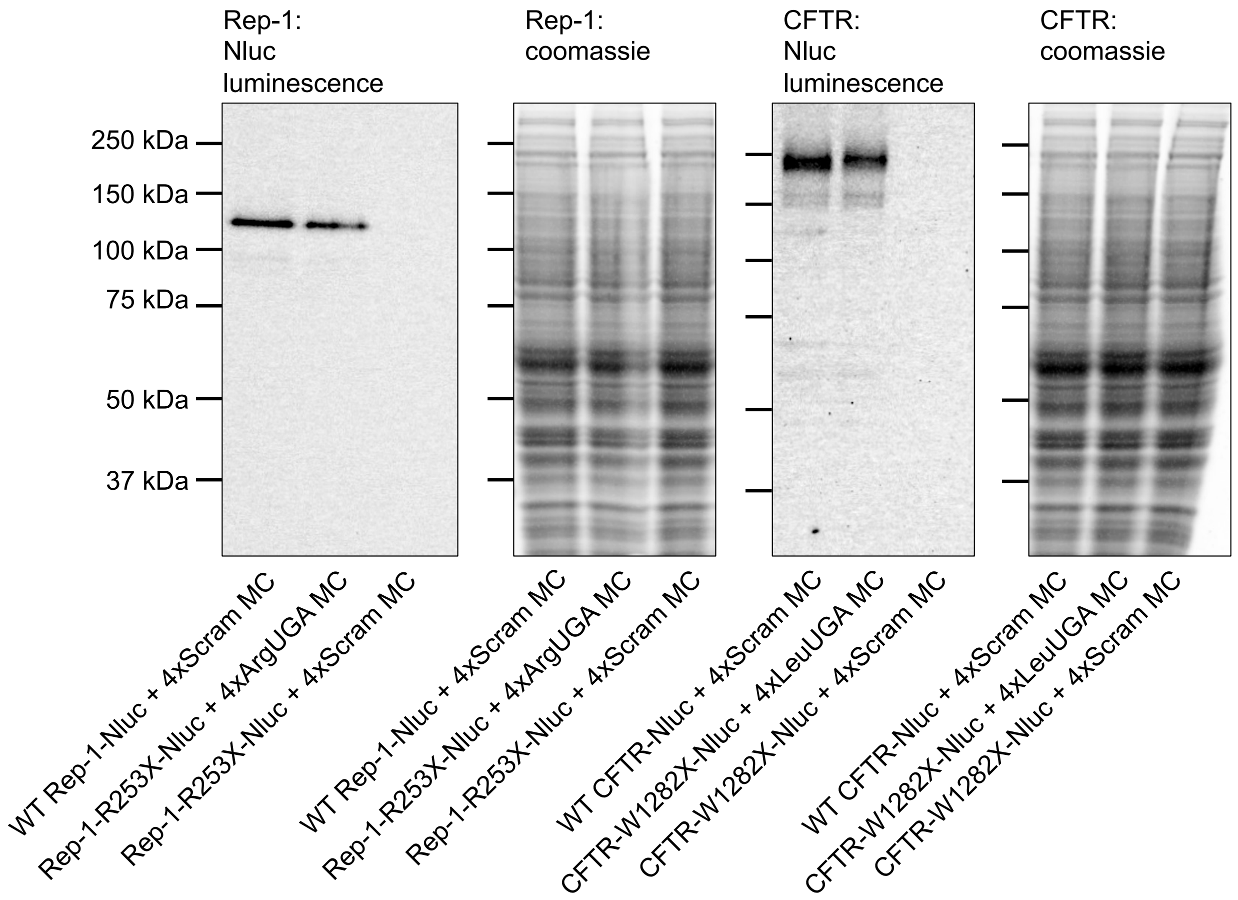


**Supplemental Figure 8. Full SDS-PAGE gel images of gels shown in Figure 2C and 2D.**

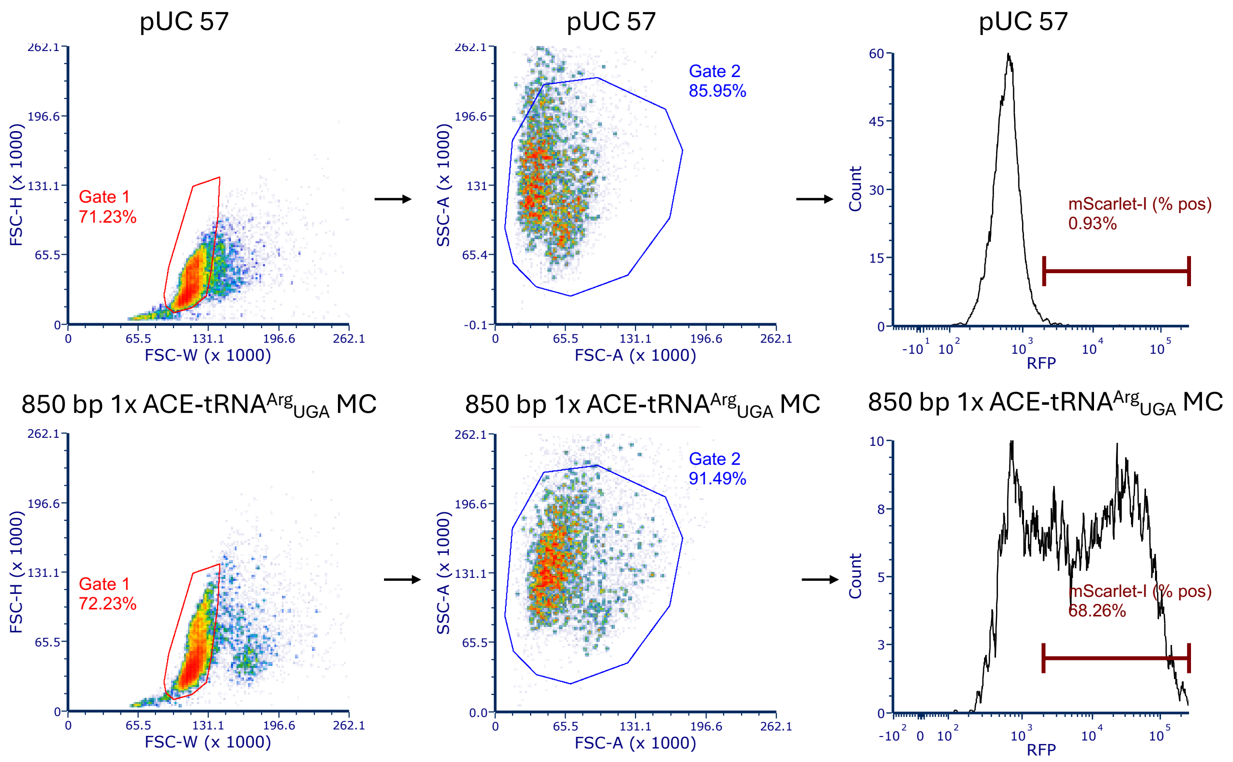


**Supplementary Figure 9. Histograms and gates for SGG flow data.** Flow cytometry data were analyzed in FCS Express 7 software, with gating for single cells accomplished using forward and side scatter, area, height, and width parameters. A gate for mScarlet-I (RFP on axis label) positive cells was established such that ~1% of events were positive in the cells treated with the negative control vector pUC 57. The percent positive population for mScarlet-I fluorescence in the ACE-tRNA minivector and pDNA treated cells was calculated based gate statistical analysis.

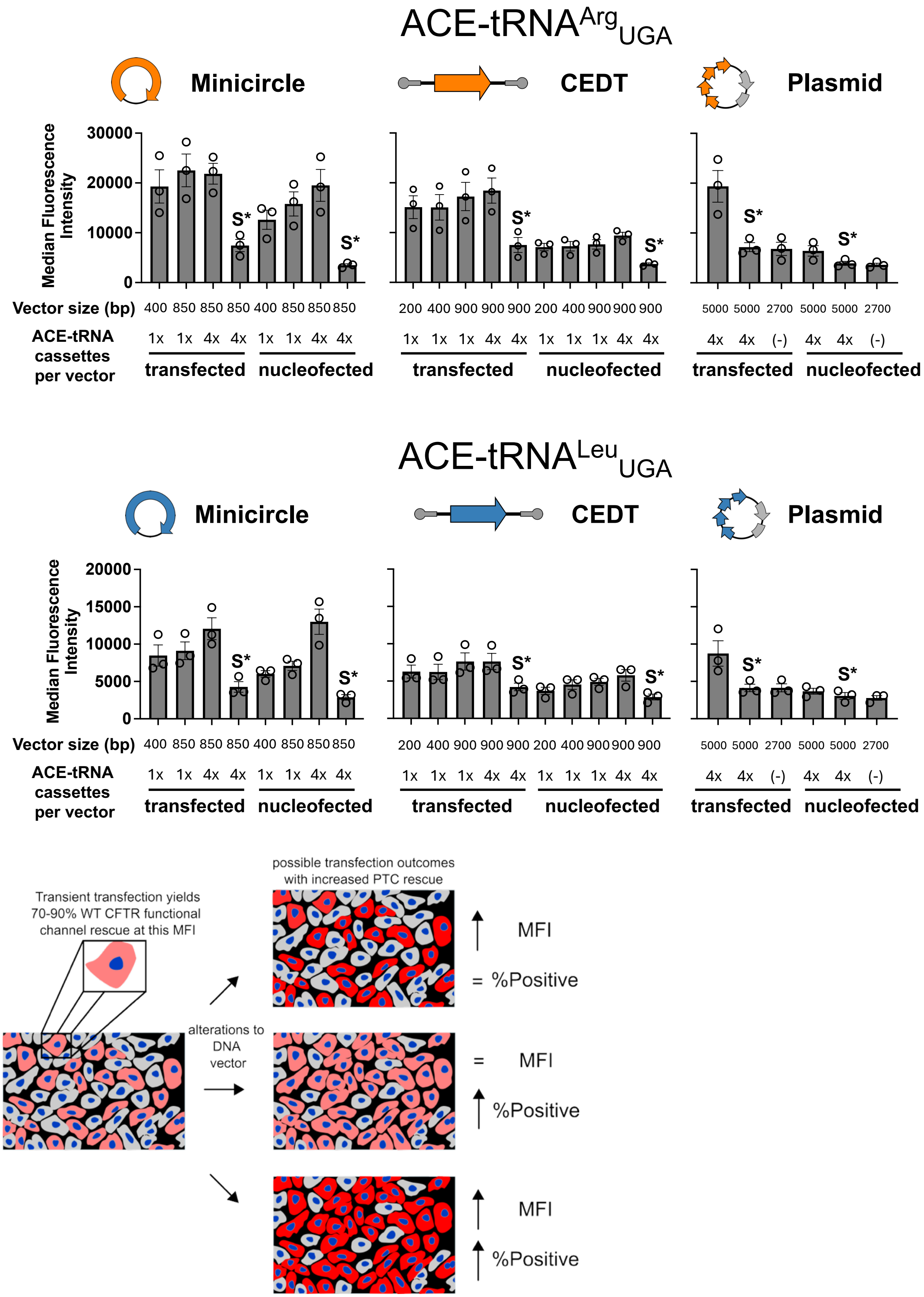
ACE-tRNA$^{Arg}_{UGA}$
Minicircle
CEDT
Plasmid
Median Fluorescence Intensity
30000
20000
10000
0
S*
Vector size (bp)
400 850 850 850 400 850 850 850
200 400 900 900 900 200 400 900 900 900
5000 5000 2700 5000 5000 2700
ACE-tRNA cassettes per vector
1x 1x 4x 4x 1x 1x 4x 4x
1x 1x 1x 4x 4x 1x 1x 1x 4x 4x
4x 4x (-) 4x 4x (-)
transfected
nucleofected
ACE-tRNA$^{Leu}_{UGA}$
Minicircle
CEDT
Plasmid
Median Fluorescence Intensity
20000
15000
10000
5000
0
S*
Vector size (bp)
400 850 850 850 400 850 850 850
200 400 900 900 900 200 400 900 900 900
5000 5000 2700 5000 5000 2700
ACE-tRNA cassettes per vector
1x 1x 4x 4x 1x 1x 4x 4x
1x 1x 1x 4x 4x 1x 1x 1x 4x 4x
4x 4x (-) 4x 4x (-)
transfected
nucleofected
Transient transfection yields 70-90% WT CFTR functional channel rescue at this MFI
alterations to DNA vector
possible transfection outcomes with increased PTC rescue
MFI
= %Positive
= MFI
%Positive
MFI
%Positive

**Supplemental Figure 10. Median fluorescence intensities from flow cytometry analysis of 16HBE14o- reporter cells transfected and nucleofected with ACE-$tRNA^{Arg}_{UGA}$ (A) or ACE-$tRNA^{Leu}_{UGA}$ (B) minivectors.** Bars represent the means of three independent transfections/nucleofections with error bars representing the S.E.M. (**C**) Alterations to DNA vectors can result in several different impacts on transfection/nucleofection outcomes. Flow cytometry reports on both the percentage of cells above a certain threshold of background fluorescence (% positive) and the aggregate fluorescence level of the population reported as the median fluorescence intensity (MFI). Depending on the influence of alterations to the physicochemical properties of DNA vectors, either the % positive, MFI, or both can increase in the cell population. In the case of transient transfection of a plasmid containing 4 copies of an ACE-tRNA expression cassette, we have previously demonstrated 70-90% rescue of CFTR channel function in 16HBEge cell lines (6). Given this previous finding, we would not necessarily seek to increase the MFI based on alterations to the DNA vectors, as the previous level of rescue was sufficient for near-WT levels of PTC rescue. Instead, we would seek to increase the % of the cell population positive for PTC rescue, as this would allow for more consistent PTC rescue across the entire PTC-affected cell population in a tissue of interest. Here we show that while the % positive increases, MFI is grossly unaffected by conversion of the ACE-tRNA from pDNA to a minivector.

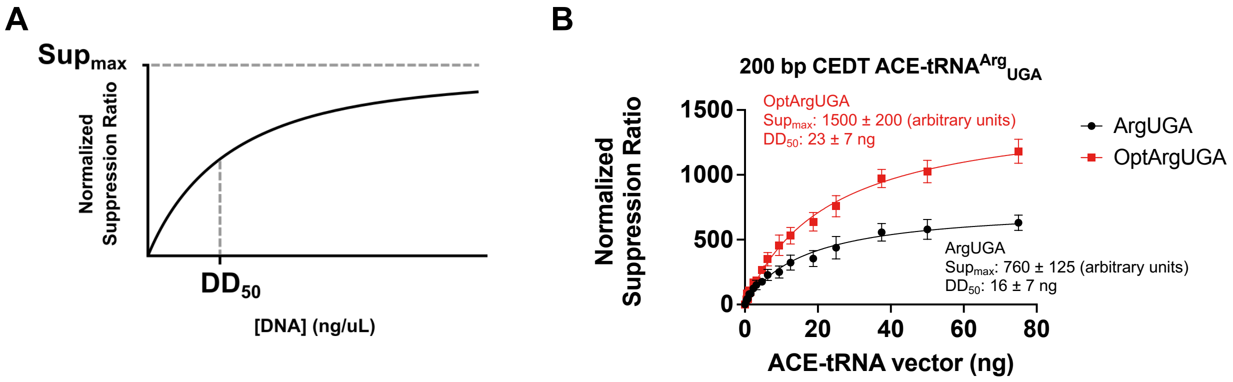


**Supplemental Figure 11. Model to fit DNA dependence of ACE-tRNA nonsense suppressor efficiency when encoded as minivectors.** (**A**) This diagram outlines the parameters of the hyperbolic model to fit the DNA concentration dependence of ACE-tRNA nonsense suppression efficiency. The model is defined by the equation Normalized Suppression Ratio = $(Sup_{max} * [DNA])/(DD_{50} + [DNA])$ where $Sup_{max}$ corresponds to the maximal level of nonsense suppression displayed by the ACE-tRNA and $DD_{50}$ (Delivered DNA) corresponds to the concentration of DNA (ng/μL) required for a half-maximal nonsense suppression response as described previously (4). (**B**) Fits of the nonsense suppressor efficiency model to both OptACE-tRNA$^{Arg}_{UGA}$ and ACE-tRNA$^{Arg}_{UGA}$ encoded as 200 bp CEDT when co-transfected into the HEK293T cell line along with the dual luciferase reporter as outlined in Fig. 2A.

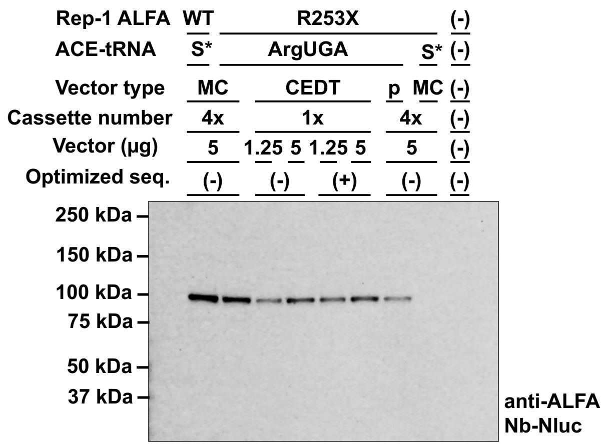


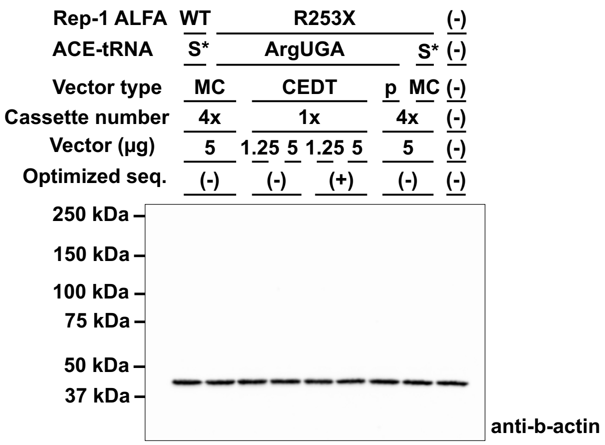


**Supplemental Figure 12. Full blot images of blots shown in Figure 4C.**

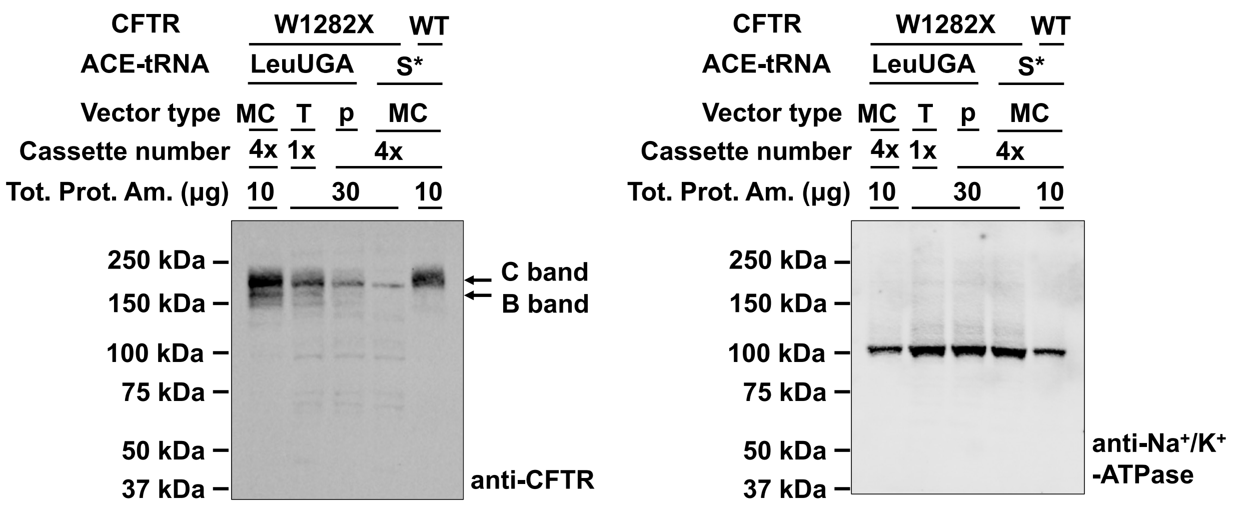


**Supplemental Figure 13. Full blot images of blots shown in Figure 5C.**

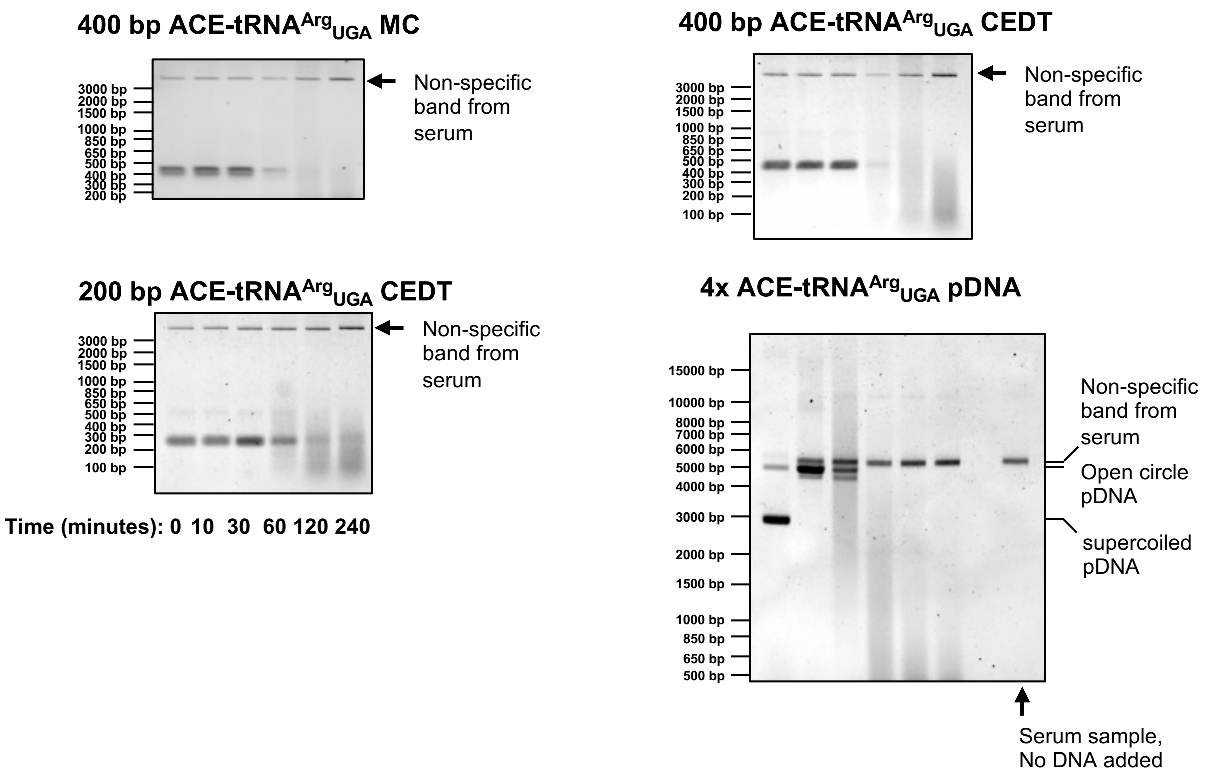


**Supplemental Figure 14. Full agarose gel images of gels shown in Figure 6C.**

| Common components of ACE-tRNA expression cassette:<br>55 bp 5' leader (derived from human tRNA gene Tyr-GTA-5-1)<br>***ACE-tRNA***<br>***Anticodon Loop***<br>4 bp 3' trailer<br>RNA Pol. III terminator | | |
|---|---|---|
| **ACE-tRNA vector name** | **ACE-tRNA vector sequence** | **Percent of vector composed of ACE-tRNA** |
| ACE-tRNA$^{Arg}_{UGA}$ (tRNA-Arg-TCT-3-1→TGA) expression cassette: | AGCGCTCCGGTTTTTCTGTGCTGAACCTCAGGGGACGCCGACACACGTA<br>CACGTC***GGCTCTGTGGCGCAATGGATAGCGCATTGGACTTCAAATTCAA***<br>***AGGTTGTGGGTTCGAGTCCCACCAGAGTCG***GTCCTTTTTTTT | N/A |
| ACE-tRNA$^{Leu}_{UGA}$ (tRNA-Leu-TAA-3-1→TGA) expression cassette: | AGCGCTCCGGTTTTTCTGTGCTGAACCTCAGGGGACGCCGACACACGTA<br>CACGTC***ACCAGAATGGCCGAGTGGTTAAGGCGTTGGACTTCAGATCCA***<br>***ATGGATTCATATCCGCGTGGGTTCGAACCCCACTTCTGGTA***GTCCTTTTTT<br>TT | N/A |
| ACE-tRNA$^{Scram}$ expression cassette: | AGCGCTCCGGTTTTTCTGTGCTGAACCTCAGGGGACGCCGACACACGTA<br>CACGTC***GGTTGGAACCTATATGACAATCCGGAAGCTGTTCGTCTAGATT***<br>***CGACGAGCCGGAGTACGCTGCCGACGTAGTTCAGTTCTAC***<br>GTCCTTTTTTTT | N/A |
| 200 bp ACE-tRNA$^{Arg}_{UGA}$ minicircle | CTTGATGGAGCAGTTATACTCTAAGCGCTCCGGTTTTTCTGTGCTGAACC<br>TCAGGGGACGCCGACACACGTACACGTC***GGCTCTGTGGCGCAATGGAT***<br>***AGCGCATTGGACTTCAAATTCAAAGGTTGTGGGTTCGAGTCCCACCAGA***<br>***GTCG***GTCCTTTTTTTTGATCCAGCAGGGTGGATGCCAGTCTGTGATTCTAA<br>GGG | 70% |
| 400 bp ACE-tRNA$^{Arg}_{UGA}$ minicircle | CAGCAGAGCTTGAACTGACTAGTTGGACACCCCACCTGAGGAGAGAGGC<br>ATAGTTATAGGATGGCCCACTACCCAAGACCCACCCTGAATGTTTTCTCTT<br>ATCTTGATGGAGCAGTTATACTCTAAGCGCTCCGGTTTTTCTGTGCTGAAC<br>CTCAGGGGACGCCGACACACGTACACGTC***GGCTCTGTGGCGCAATGGA***<br>***TAGCGCATTGGACTTCAAATTCAAAGGTTGTGGGTTCGAGTCCCACCAG***<br>***AGTCG***GTCCTTTTTTTTGATCCAGCAGGGTGGATGCCAGTCTGTGATTCTA<br>AGGGCTTAAACCTTATAGTTAGGGGAAATGCTTGGGGTACTTGAACACAA<br>ACATGTCCCAAGCTGGGGGTCCTAGCCAAAGATGAGCCTTTCTAACTAGC | 35% |
| 850 bp 1x ACE-tRNA$^{Arg}_{UGA}$ minicircle (phiC31 attR shown in lowercase) | ccccaactggggtaacctttgggctccccgggcgcgTGCCGGGGGGTGCATGCAACCCCC<br>CTAACCTCCACTGAAACCTTGATGGGAAGTCCAATTTTAGGGTAGGAACC<br>TGATTTGGGGGTCCTTAAGCACCATGGGGACAATGCTGACCACTATGGG<br>GTGGGTTGGGGACCTGGACCCCAATTGATTTCCTGTATTGCTAGAAATGC<br>TGATGCTAAGCTGGGCCCATTAGTTCTGACAGACCTGGGGCCCTAGAAG<br>TCAGAGCTTGAACTGACTAGTTGGACACCCCACCTGAGGAGAGAGGCAT<br>AGTTATAGGATGGCCCACTACCCAAGACCCACCCTGAATGTTTTCTCTTAT<br>CTTGATGGAGCAGTTATACTCTAGACTCACTATAGAGCGCTCCGGTTTTTC<br>TGTGCTGAACCTCAGGGGACGCCGACACACGTACACGTC***GGCTCTGTGG*** | 16% |

| | | |
|---|---|---|
| | CGCAATGGATAGCGCATTGGACTTCAAATTCAAAGGTTGTGGGTTCGAG<br>TCCCACCAGAGTCGGTCCTTTTTTTGCTTTAGTGAGGGTTAATTCAGCAT<br>GATCCAGCAGGGTGGATGCCAGTCTGTGATTCTAAGGGCTTAAACCTTAT<br>AGTTAGGGGAAATGCTTGGGGTACTTGAACACAAACATGTCCCAAGCTGG<br>GGGTCCTAGCCAAAGATGAGCCTTTCTAACTAGCCTCAGGCCTTAGTTTG<br>GTAATAAAAAAGCAGGCCAATCTATTTCTGCCATGGGTGATGAGTGTCAT<br>CCCCTATTGATTGGACAGGGGTTAAGTGGTGAGCCATGCATCCAGATAGT<br>GCTAGATTATAAGGACCCTATAGAACAGGTCTGTAAATTTACATAGTTCCA<br>TTTATTAAACTGGAACATAGAATTGAGGCAA | |
| 850 bp 4x ACE-$tRNA^{Arg}_{UGA}$ minicircle (phiC31 attR shown in lowercase) | ccccaactggggtaacctttgggctccccgggcgcgTGCCGATGGAGCAGTTATACTCTAG<br>ACTCACTATAGAGCGCTCCGGTTTTTCTGTGCTGAACCTCAGGGGACGCC<br>GACACACGTACACGTCGGCTCTGTGGCGCAATGGATAGCGCATTGGACT<br>TCAAATTCAAAGGTTGTGGGTTCGAGTCCCACCAGAGTCGGTCCTTTTTT<br>TGCTTTAGTGAGGGTTAATTCAGCATGATCCactaGATGGAGCAGTTATACT<br>CTAGACTCACTATAGAGCGCTCCGGTTTTTCTGTGCTGAACCTCAGGGGA<br>CGCCGACACACGTACACGTCGGCTCTGTGGCGCAATGGATAGCGCATTG<br>GACTTCAAATTCAAAGGTTGTGGGTTCGAGTCCCACCAGAGTCGGTCCT<br>TTTTTTGCTTTAGTGAGGGTTAATTCAGCATGATCCttacGATGGAGCAGTT<br>ATACTCTAGACTCACTATAGAGCGCTCCGGTTTTTCTGTGCTGAACCTCA<br>GGGGACGCCGACACACGTACACGTCGGCTCTGTGGCGCAATGGATAGC<br>GCATTGGACTTCAAATTCAAAGGTTGTGGGTTCGAGTCCCACCAGAGTC<br>GGTCCTTTTTTTGCTTTAGTGAGGGTTAATTCAGCATGATCCcagaGATGGA<br>GCAGTTATACTCTAGACTCACTATAGAGCGCTCCGGTTTTTCTGTGCTGAA<br>CCTCAGGGGACGCCGACACACGTACACGTCGGCTCTGTGGCGCAATGG<br>ATAGCGCATTGGACTTCAAATTCAAAGGTTGTGGGTTCGAGTCCCACCA<br>GAGTCGGTCCTTTTTTTGCTTTAGTGAGGGTTAATTCAGCATGATCCGCAA | 65% |
| 850 bp 4x ACE-$tRNA^{Scram}$ minicircle (phiC31 attR shown in lowercase) | ccccaactggggtaacctttgggctccccgggcgcgTGCCGATGGAGCAGTTATACTCTAG<br>ACTCACTATAGAGCGCTCCGGTTTTTCTGTGCTGAACCTCAGGGGACGCC<br>GACACACGTACACGTCGGTTGGAACCTATATGACAATCCGGAAGCTGTT<br>CGTCTAGATTCGACGAGCCGGAGTACGCTGCCGACGTAGTTCAGTTCTA<br>CGTCCTTTTTTTGCTTTAGTGAGGGTTAATTCAGCATGATCCactaGATGGA<br>GCAGTTATACTCTAGACTCACTATAGAGCGCTCCGGTTTTTCTGTGCTGAA<br>CCTCAGGGGACGCCGACACACGTACACGTCGGTTGGAACCTATATGACA<br>ATCCGGAAGCTGTTCGTCTAGATTCGACGAGCCGGAGTACGCTGCCGA<br>CGTAGTTCAGTTCTACGTCCTTTTTTTGCTTTAGTGAGGGTTAATTCAGCA<br>TGATCCttacGATGGAGCAGTTATACTCTAGACTCACTATAGAGCGCTCCGG<br>TTTTTCTGTGCTGAACCTCAGGGGACGCCGACACACGTACACGTCGGTTG<br>GAACCTATATGACAATCCGGAAGCTGTTCGTCTAGATTCGACGAGCCGG<br>AGTACGCTGCCGACGTAGTTCAGTTCTACGTCCTTTTTTTGCTTTAGTGA<br>GGGTTAATTCAGCATGATCCcagaGATGGAGCAGTTATACTCTAGACTCAC<br>TATAGAGCGCTCCGGTTTTTCTGTGCTGAACCTCAGGGGACGCCGACAC<br>ACGTACACGTCGGTTGGAACCTATATGACAATCCGGAAGCTGTTCGTCT<br>AGATTCGACGAGCCGGAGTACGCTGCCGACGTAGTTCAGTTCTACGTCC<br>TTTTTTTGCTTTAGTGAGGGTTAATTCAGCATGATCCGCAA | 65% |
| 200 bp ACE-$tRNA^{Arg}_{UGA}$ CEDT (TelL highlighted green, TelR yellow) | GCGTATAATGGACTATTGTGTGCTGATATTGATGGAGCAGTTATACTCTA<br>AGCGCTCCGGTTTTTCTGTGCTGAACCTCAGGGGACGCCGACACACGTA<br>CACGTCGGCTCTGTGGCGCAATGGATAGCGCATTGGACTTCAAATTCAA<br>AGGTTGTGGGTTCGAGTCCCACCAGAGTCGGTCCTTTTTTTGATCCAGCA<br>GGGTGGATGCCAGTCTGTGATTCTAAGGTATCAGCACACAATTGCCCAT<br>TAT | 55% |
| 400 bp ACE-$tRNA^{Arg}_{UGA}$ CEDT (TelL highlighted green, TelR yellow) | GCGTATAATGGACTATTGTGTGCTCCTGCTTGAACTGACTAGTTGGACAC<br>CCCACCTGAGGAGAGAGGCATAGTTATAGGATGGCCCACTACCCAAGAC<br>CCACCCTGAATGTTTTCTCTTATCTTGATGGAGCAGTTATACTCTAAGCGC<br>TCCGGTTTTTCTGTGCTGAACCTCAGGGGACGCCGACACACGTACACGT<br>CGGCTCTGTGGCGCAATGGATAGCGCATTGGACTTCAAATTCAAAGGTT<br>GTGGGTTCGAGTCCCACCAGAGTCGGTCCTTTTTTTGATCCAGCAGGGT<br>GGATGCCAGTCTGTGATTCTAAGGGCTTAAACCTTATAGTTAGGGGAAAT<br>GCTTGGGGTACTTGAACACAAACATGTCCCAAGCTGGGGGTCCTAGCCA<br>AAGATGAGCCTTTCTAACTAGCTATCTATCAGCACACAATTGCCCATTAT | 31% |
| 900 bp 1xACE-$tRNA^{Arg}_{UGA}$ CEDT (TelL highlighted green, TelR yellow) | GCGTATAATGGACTATTGTGTGCTCTGGGGGCATCATATAGCTGCATCTT<br>CTCTGGGGGGGTGCATGCAACCCCCCTAACCTCCACTGAAACCTTGATG<br>GGAAGTCCAATTTTAGGGTAGGAACCTGATTTGGGGGTCCTTAAGCACCA<br>TGGGGACAATGCTGACCACTATGGGGTGGGTTGGGGACCTGGACCCCAA<br>TTGATTTCCTGTATTGCTAGAAATGCTGATGCTAAGCTGGGCCCATTAGTT<br>CTGACAGACCTGGGGCCCTAGAAGTCAGAGCTTGAACTGACTAGTTGGA<br>CACCCCACCTGAGGAGAGAGGCATAGTTATAGGATGGCCCACTACCCAA<br>GACCCACCCTGAATGTTTTCTCTTATCTTGATGGAGCAGTTATACTCTAAG<br>CGCTCCGGTTTTTCTGTGCTGAACCTCAGGGGACGCCGACACACGTACA<br>CGTCGGCTCTGTGGCGCAATGGATAGCGCATTGGACTTCAAATTCAAAG<br>GTTGTGGGTTCGAGTCCCACCAGAGTCGGTCCTTTTTTTGATCCAGCAG | 15% |

| | | |
|---|---|---|
| | GGTGGATGCCAGTCTGTGATTCTAAGGGCTTAAACCTTATAGTTAGGGGA<br>AATGCTTGGGGTACTTGAACACAAACATGTCCCAAGCTGGGGGTCCTAGC<br>CAAAGATGAGCCTTTCTAACTAGCCTCAGGCCTTAGTTTGGTAATAAAAA<br>GCAGGCCAATCTATTTCTGCCATGGGTGATGAGTGTCATCCCCTATTGAT<br>TGGACAGGGGTTAAGTGGTGAGCCATGCATCCAGATAGTGCTAGATTATA<br>AGGACCCTATAGAACAGGTCTGTAAATTTACATAGTTCCATTTATTAAACT<br>GGAACATAGAATTGAGAAATCTTAGGGTGCCATTAAAGACACCCTGCATT<br>CCTTCTGACCACAAGCTGTTATGCTAA**TATCAGCACACAATTGCCCATTA**<br>**T** | |
| 900 bp 4xACE-tRNA$^{Arg}_{UGA}$ CEDT (TelL highlighted green, TelR yellow) | **GCGTATAATGGACTATTGTGTGCT**cctgAGGGCTTGATGGAGCAGTTATAC<br>TCTAGACTCACTATAGAGCGCTCCGGTTTTTCTGTGCTGAACCTCAGGGG<br>ACGCCGACACACGTACACGTC***GGTTGGAACCTATATGACAATCCGGAAG***<br>***CTGTTCGTCTAGATTCGACGAGCCGGAGTACGCTGCCGACGTAGTTCAG***<br>***TTCTAC***GTCCTTTTTTTGCTTTAGTGAGGGTTAATTCAGCATGATCCactaG<br>ATGGAGCAGTTATACTCTAGACTCACTATAGAGCGCTCCGGTTTTTCTGT<br>GCTGAACCTCAGGGGACGCCGACACACGTACACGTC***GGTTGGAACCTAT***<br>***ATGACAATCCGGAAGCTGTTCGTCTAGATTCGACGAGCCGGAGTACGCT***<br>***GCCGACGTAGTTCAGTTCTAC***GTCCTTTTTTTGCTTTAGTGAGGGTTAATT<br>CAGCATGATCCttacGATGGAGCAGTTATACTCTAGACTCACTATAGAGCGC<br>TCCGGTTTTTCTGTGCTGAACCTCAGGGGACGCCGACACACGTACACGT<br>C***GGTTGGAACCTATATGACAATCCGGAAGCTGTTCGTCTAGATTCGACG***<br>***AGCCGGAGTACGCTGCCGACGTAGTTCAGTTCTAC***GTCCTTTTTTTGCTT<br>TAGTGAGGGTTAATTCAGCATGATCCcagaGATGGAGCAGTTATACTCTAG<br>ACTCACTATAGAGCGCTCCGGTTTTTCTGTGCTGAACCTCAGGGGACGCC<br>GACACACGTACACGTC***GGTTGGAACCTATATGACAATCCGGAAGCTGTT***<br>***CGTCTAGATTCGACGAGCCGGAGTACGCTGCCGACGTAGTTCAGTTCTA***<br>***C***GTCCTTTTTTTGCTTTAGTGAGGGTTAATTCAGCATGATCCAGCAGGGT<br>GGATGCCAGTCTGTGATTCTAAGGG**TATCAGCACACAATTGCCCATTAT** | 61% |
| 900 bp 4xACE-tRNA$^{Scram}$ CEDT (TelL highlighted green, TelR yellow) | **GCGTATAATGGACTATTGTGTGCT**cctgAGGGCTTGATGGAGCAGTTATAC<br>TCTAGACTCACTATAGAGCGCTCCGGTTTTTCTGTGCTGAACCTCAGGGG<br>ACGCCGACACACGTACACGTC***GGTTGGAACCTATATGACAATCCGGAAG***<br>***CTGTTCGTCTAGATTCGACGAGCCGGAGTACGCTGCCGACGTAGTTCAG***<br>***TTCTAC***GTCCTTTTTTTGCTTTAGTGAGGGTTAATTCAGCATGATCCactaG<br>ATGGAGCAGTTATACTCTAGACTCACTATAGAGCGCTCCGGTTTTTCTGT<br>GCTGAACCTCAGGGGACGCCGACACACGTACACGTC***GGTTGGAACCTAT***<br>***ATGACAATCCGGAAGCTGTTCGTCTAGATTCGACGAGCCGGAGTACGCT***<br>***GCCGACGTAGTTCAGTTCTAC***GTCCTTTTTTTGCTTTAGTGAGGGTTAATT<br>CAGCATGATCCttacGATGGAGCAGTTATACTCTAGACTCACTATAGAGCGC<br>TCCGGTTTTTCTGTGCTGAACCTCAGGGGACGCCGACACACGTACACGT<br>C***GGTTGGAACCTATATGACAATCCGGAAGCTGTTCGTCTAGATTCGACG***<br>***AGCCGGAGTACGCTGCCGACGTAGTTCAGTTCTAC***GTCCTTTTTTTGCTT<br>TAGTGAGGGTTAATTCAGCATGATCCcagaGATGGAGCAGTTATACTCTAG<br>ACTCACTATAGAGCGCTCCGGTTTTTCTGTGCTGAACCTCAGGGGACGCC<br>GACACACGTACACGTC***GGTTGGAACCTATATGACAATCCGGAAGCTGTT***<br>***CGTCTAGATTCGACGAGCCGGAGTACGCTGCCGACGTAGTTCAGTTCTA***<br>***C***GTCCTTTTTTTGCTTTAGTGAGGGTTAATTCAGCATGATCCAGCAGGGT<br>GGATGCCAGTCTGTGATTCTAAGGG**TATCAGCACACAATTGCCCATTAT** | 63% |
| 4xACE-tRNA$^{Arg}_{UGA}$ pMC insert (whole vector is 5 kb, phiC31 attB highlighted blue, phiC31 attP39 highlighted purple) | GTGCCAGGGCGTGCCCTTGGGCTCCCCGGGCGCGtgccGATGGAGCAGT<br>TATACTCTAGACTCACTATAGAGCGCTCCGGTTTTTCTGTGCTGAACCTCA<br>GGGGACGCCGACACACGTACACGTC***GGTTGGAACCTATATGACAATCCG***<br>***GAAGCTGTTCGTCTAGATTCGACGAGCCGGAGTACGCTGCCGACGTAG***<br>***TTCAGTTCTAC***GTCCTTTTTTTGCTTTAGTGAGGGTTAATTCAGCATGATC<br>CactaGATGGAGCAGTTATACTCTAGACTCACTATAGAGCGCTCCGGTTTTT<br>CTGTGCTGAACCTCAGGGGACGCCGACACACGTACACGTC***GGTTGGAAC***<br>***CTATATGACAATCCGGAAGCTGTTCGTCTAGATTCGACGAGCCGGAGTA***<br>***CGCTGCCGACGTAGTTCAGTTCTAC***GTCCTTTTTTTGCTTTAGTGAGGGT<br>TAATTCAGCATGATCCttacGATGGAGCAGTTATACTCTAGACTCACTATAG<br>AGCGCTCCGGTTTTTCTGTGCTGAACCTCAGGGGACGCCGACACACGTA<br>CACGTC***GGTTGGAACCTATATGACAATCCGGAAGCTGTTCGTCTAGATT***<br>***CGACGAGCCGGAGTACGCTGCCGACGTAGTTCAGTTCTAC***GTCCTTTTT<br>TTGCTTTAGTGAGGGTTAATTCAGCATGATCCcagaGATGGAGCAGTTATA<br>CTCTAGACTCACTATAGAGCGCTCCGGTTTTTCTGTGCTGAACCTCAGGG<br>GACGCCGACACACGTACACGTC***GGTTGGAACCTATATGACAATCCGGAA***<br>***GCTGTTCGTCTAGATTCGACGAGCCGGAGTACGCTGCCGACGTAGTTCA***<br>***GTTCTAC***GTCCTTTTTTTGCTTTAGTGAGGGTTAATTCAGCATGATCCgcaa<br>GCCCCAACTGGGGTAACCTTTGAGTTCTCTCAGTTGGGGG | 11% |
| 4xACE-tRNA$^{Scram}$ pMC insert (whole | GTGCCAGGGCGTGCCCTTGGGCTCCCCGGGCGCGtgccGATGGAGCAGT<br>TATACTCTAGACTCACTATAGAGCGCTCCGGTTTTTCTGTGCTGAACCTCA<br>GGGGACGCCGACACACGTACACGTC***GGTTGGAACCTATATGACAATCCG***<br>***GAAGCTGTTCGTCTAGATTCGACGAGCCGGAGTACGCTGCCGACGTAG*** | 12% |

| | | |
|---|---|---|
| vector is 5 kb, phiC31 attB highlighted blue, phiC31 attP39 highlighted purple) | TTCAGTTCTACGTCCTTTTTTTGCTTTAGTGAGGGTTAATTCAGCATGATC<br>CactaGATGGAGCAGTTATACTCTAGACTCACTATAGAGCGCTCCGGTTTTT<br>CTGTGCTGAACCTCAGGGGACGCCGACACACGTACACGTCGGTTGGAAC<br>CTATATGACAATCCGGAAGCTGTTCGTCTAGATTCGACGAGCCGGAGTA<br>CGCTGCCGACGTAGTTCAGTTCTACGTCCTTTTTTTGCTTTAGTGAGGGT<br>TAATTCAGCATGATCCttacGATGGAGCAGTTATACTCTAGACTCACTATAG<br>AGCGCTCCGGTTTTTCTGTGCTGAACCTCAGGGGACGCCGACACACGTA<br>CACGTCGGTTGGAACCTATATGACAATCCGGAAGCTGTTCGTCTAGATT<br>CGACGAGCCGGAGTACGCTGCCGACGTAGTTCAGTTCTACGTCCTTTTT<br>TTGCTTTAGTGAGGGTTAATTCAGCATGATCCcagaGATGGAGCAGTTATA<br>CTCTAGACTCACTATAGAGCGCTCCGGTTTTTCTGTGCTGAACCTCAGGG<br>GACGCCGACACACGTACACGTCGGTTGGAACCTATATGACAATCCGGAA<br>GCTGTTCGTCTAGATTCGACGAGCCGGAGTACGCTGCCGACGTAGTTCA<br>GTTCTACGTCCTTTTTTTGCTTTAGTGAGGGTTAATTCAGCATGATCCgcaa<br>GCCCCAACTGGGGTAACCTTTGAGTTCTCTCAGTTGGGGG | |
| 200 bp ACE-tRNA$^{Leu}_{UGA}$ minicircle | CTTGATGGAGCAGTTATACTCTAAGCGCTCCGGTTTTTCTGTGCTGAACC<br>TCAGGGGACGCCGACACACGTACACGTCACCAGAATGGCCGAGTGGTT<br>AAGGCGTTGGACTTCAGATCCAATGGATTCATATCCGCGTGGGTTCGAA<br>CCCCACTTCTGGTAGTCCTTTTTTTGATCCAGCAGGGTGGATGCCAGTCT<br>GTGATTCTAAGGG | 71% |
| 400 bp ACE-tRNA$^{Leu}_{UGA}$ minicircle | CAGCAGAGCTTGAACTGACTAGTTGGACACCCCACCTGAGGAGAGAGGC<br>ATAGTTATAGGATGGCCCACTACCCAAGACCCACCCTGAATGTTTTCTCTT<br>ATCTTGATGGAGCAGTTATACTCTAAGCGCTCCGGTTTTTCTGTGCTGAAC<br>CTCAGGGGACGCCGACACACGTACACGTCACCAGAATGGCCGAGTGGT<br>TAAGGCGTTGGACTTCAGATCCAATGGATTCATATCCGCGTGGGTTCGA<br>ACCCCACTTCTGGTAGTCCTTTTTTTGATCCAGCAGGGTGGATGCCAGTC<br>TGTGATTCTAAGGGCTTAAACCTTATAGTTAGGGGAAATGCTTGGGGTAC<br>TTGAACACAAACATGTCCCAAGCTGGGGGTCCTAGCCAAAGATGAGCCTT<br>TCTAACTAGC | 37% |
| 850 bp 1x ACE-tRNA$^{Leu}_{UGA}$ minicircle (phiC31 attR shown in lowercase) | ccccaactggggtaacctttgggctccccgggcgcgTGCCGGGGGGTGCATGCAACCCCC<br>CTAACCTCCACTGAAACCTTGATGGGAAGTCCAATTTTAGGGTAGGAACC<br>TGATTTGGGGGTCCTTAAGCACCATGGGGACAATGCTGACCACTATGGG<br>GTGGGTTGGGGACCTGGACCCCAATTGATTTCCTGTATTGCTAGAAATGC<br>TGATGCTAAGCTGGGCCCATTAGTTCTGACAGACCTGGGGCCCTAGAAG<br>TCAGAGCTTGAACTGACTAGTTGGACACCCCACCTGAGGAGAGAGGCAT<br>AGTTATAGGATGGCCCACTACCCAAGACCCACCCTGAATGTTTTCTCTTAT<br>CTTGATGGAGCAGTTATACTCTAGACTCACTATAGAGCGCTCCGGTTTTTC<br>TGTGCTGAACCTCAGGGGACGCCGACACACGTACACGTCACCAGAATGG<br>CCGAGTGGTTAAGGCGTTGGACTTCAGATCCAATGGATTCATATCCGCG<br>TGGGTTCGAACCCCACTTCTGGTAGTCCTTTTTTTGCTTTAGTGAGGGTT<br>AATTCAGCATGATCCAGCAGGGTGGATGCCAGTCTGTGATTCTAAGGGCT<br>TAAACCTTATAGTTAGGGGAAATGCTTGGGGTACTTGAACACAAACATGT<br>CCCAAGCTGGGGGTCCTAGCCAAAGATGAGCCTTTCTAACTAGCCTCAG<br>GCCTTAGTTTGGTAATAAAAAGCAGGCCAATCTATTTCTGCCATGGGTG<br>ATGAGTGTCATCCCCTATTGATTGGACAGGGGTTAAGTGGTGAGCCATGC<br>ATCCAGATAGTGCTAGATTATAAGGACCCTATAGAACAGGTCTGTAAATTT<br>ACATAGTTCCATTTATTAAACTGGAACATAGAATTGAGGCAA | 17% |
| 850 bp 4x ACE-tRNA$^{Leu}_{UGA}$ minicircle (phiC31 attR shown in lowercase) | ccccaactggggtaacctttgggctccccgggcgcgTGCCGATGGAGCAGTTATACTCTAG<br>ACTCACTATAGAGCGCTCCGGTTTTTCTGTGCTGAACCTCAGGGGACGCC<br>GACACACGTACACGTCACCAGAATGGCCGAGTGGTTAAGGCGTTGGACT<br>TCAGATCCAATGGATTCATATCCGCGTGGGTTCGAACCCCACTTCTGGT<br>AGTCCTTTTTTTGCTTTAGTGAGGGTTAATTCAGCATGATCCactaGATGGA<br>GCAGTTATACTCTAGACTCACTATAGAGCGCTCCGGTTTTTCTGTGCTGAA<br>CCTCAGGGGACGCCGACACACGTACACGTCACCAGAATGGCCGAGTGG<br>TTAAGGCGTTGGACTTCAGATCCAATGGATTCATATCCGCGTGGGTTCG<br>AACCCCACTTCTGGTAGTCCTTTTTTTGCTTTAGTGAGGGTTAATTCAGCA<br>TGATCCttacGATGGAGCAGTTATACTCTAGACTCACTATAGAGCGCTCCGG<br>TTTTTCTGTGCTGAACCTCAGGGGACGCCGACACACGTACACGTCACCA<br>GAATGGCCGAGTGGTTAAGGCGTTGGACTTCAGATCCAATGGATTCATA<br>TCCGCGTGGGTTCGAACCCCACTTCTGGTAGTCCTTTTTTTGCTTTAGTG<br>AGGGTTAATTCAGCATGATCCcagaGATGGAGCAGTTATACTCTAGACTCA<br>CTATAGAGCGCTCCGGTTTTTCTGTGCTGAACCTCAGGGGACGCCGACA<br>CACGTACACGTCACCAGAATGGCCGAGTGGTTAAGGCGTTGGACTTCAG<br>ATCCAATGGATTCATATCCGCGTGGGTTCGAACCCCACTTCTGGTAGTC<br>CTTTTTTTGCTTTAGTGAGGGTTAATTCAGCATGATCCGCAA | 66% |
| 200 bp ACE-tRNA$^{Leu}_{UGA}$ CEDT | GCGTATAATGGACTATTGTGTGCTGATATTGATGGAGCAGTTATACTCTA<br>AGCGCTCCGGTTTTTCTGTGCTGAACCTCAGGGGACGCCGACACACGTA<br>CACGTCACCAGAATGGCCGAGTGGTTAAGGCGTTGGACTTCAGATCCA<br>ATGGATTCATATCCGCGTGGGTTCGAACCCCACTTCTGGTAGTCCTTTTT | 57% |

| | | |
|---|---|---|
| (TelL highlighted green, TelR yellow) | TTGATCCAGCAGGGTGGATGCCAGTCTGTGATTCTAAGG**TATCAGCACA**<br>**CAATTGCCCATTAT** | |
| 400 bp ACE-tRNA$^{Leu}_{UGA}$ CEDT (TelL highlighted green, TelR yellow) | **GCGTATAATGGACTATTGTGTGCT**CCTGCTTGAACTGACTAGTTGGACAC<br>CCCACCTGAGGAGAGAGGCATAGTTATAGGATGGCCCACTACCCAAGAC<br>CCACCCTGAATGTTTTCTCTTATCTTGATGGAGCAGTTATACTCTAAGCGC<br>TCCGGTTTTTCTGTGCTGAACCTCAGGGGACGCCGACACACGTACACGT<br>C***ACCAGAATGGCCGAGTGGTTAAGGCGTTGGACTTCAGATCCAATGGA***<br>***TTCATATCCGCGTGGGTTCGAACCCCACTTCTGGTA***GTCCTTTTTTTGAT<br>CCAGCAGGGTGGATGCCAGTCTGTGATTCTAAGGGCTTAAACCTTATAGT<br>TAGGGGAAATGCTTGGGGTACTTGAACACAAACATGTCCCAAGCTGGGG<br>GTCCTAGCCAAAGATGAGCCTTTCTAACTAGCTATC**TATCAGCACACAAT**<br>**TGCCCATTAT** | 33% |
| 900 bp 1xACE-tRNA$^{Leu}_{UGA}$ CEDT (TelL highlighted green, TelR yellow) | **GCGTATAATGGACTATTGTGTGCT**CTGGGGGCATCATATAGCTGCATCTT<br>CTCTGGGGGGGTGCATGCAACCCCCCTAACCTCCACTGAAACCTTGATG<br>GGAAGTCCAATTTTAGGGTAGGAACCTGATTTGGGGGTCCTTAAGCACCA<br>TGGGGACAATGCTGACCACTATGGGGTGGGTTGGGGACCTGGACCCCAA<br>TTGATTTCCTGTATTGCTAGAAATGCTGATGCTAAGCTGGGCCCATTAGTT<br>CTGACAGACCTGGGGCCCTAGAAGTCAGAGCTTGAACTGACTAGTTGGA<br>CACCCCACCTGAGGAGAGAGGCATAGTTATAGGATGGCCCACTACCCAA<br>GACCCACCCTGAATGTTTTCTCTTATCTTGATGGAGCAGTTATACTCTAAG<br>CGCTCCGGTTTTTCTGTGCTGAACCTCAGGGGACGCCGACACACGTACA<br>CGTC***ACCAGAATGGCCGAGTGGTTAAGGCGTTGGACTTCAGATCCAAT***<br>***GGATTCATATCCGCGTGGGTTCGAACCCCACTTCTGGTA***GTCCTTTTTTT<br>GATCCAGCAGGGTGGATGCCAGTCTGTGATTCTAAGGGCTTAAACCTTAT<br>AGTTAGGGGAAATGCTTGGGGTACTTGAACACAAACATGTCCCAAGCTGG<br>GGGTCCTAGCCAAAGATGAGCCTTTCTAACTAGCCTCAGGCCTTAGTTTG<br>GTAATAAAAAAGCAGGCCAATCTATTTCTGCCATGGGTGATGAGTGTCAT<br>CCCCTATTGATTGGACAGGGGTTAAGTGGTGAGCCATGCATCCAGATAGT<br>GCTAGATTATAAGGACCCTATAGAACAGGTCTGTAAATTTACATAGTTCCA<br>TTTATTAAACTGGAACATAGAATTGAGAAATCTTAGGGTGCCATTAAAGAC<br>ACCCTGCATTCCTTCTGACCACAAGCTGTTATGCTAA**TATCAGCACACAA**<br>**TTGCCCATTAT** | 16% |
| 900 bp 4xACE-tRNA$^{Leu}_{UGA}$ CEDT (TelL highlighted green, TelR yellow) | **GCGTATAATGGACTATTGTGTGCT**cctgAGGGCTTGATGGAGCAGTTATAC<br>TCTAGACTCACTATAGAGCGCTCCGGTTTTTCTGTGCTGAACCTCAGGGG<br>ACGCCGACACACGTACACGTC***ACCAGAATGGCCGAGTGGTTAAGGCGTT***<br>***GGACTTCAGATCCAATGGATTCATATCCGCGTGGGTTCGAACCCCACTT***<br>***CTGGTA***GTCCTTTTTTTGCTTTAGTGAGGGTTAATTCAGCATGATCCactaG<br>ATGGAGCAGTTATACTCTAGACTCACTATAGAGCGCTCCGGTTTTTCTGT<br>GCTGAACCTCAGGGGACGCCGACACACGTACACGTC***ACCAGAATGGCC***<br>***GAGTGGTTAAGGCGTTGGACTTCAGATCCAATGGATTCATATCCGCGTG***<br>***GGTTCGAACCCCACTTCTGGTA***GTCCTTTTTTTGCTTTAGTGAGGGTTAAT<br>TCAGCATGATCCttacGATGGAGCAGTTATACTCTAGACTCACTATAGAGCG<br>CTCCGGTTTTTCTGTGCTGAACCTCAGGGGACGCCGACACACGTACACG<br>TC***ACCAGAATGGCCGAGTGGTTAAGGCGTTGGACTTCAGATCCAATGG***<br>***ATTCATATCCGCGTGGGTTCGAACCCCACTTCTGGTA***GTCCTTTTTTTGC<br>TTTAGTGAGGGTTAATTCAGCATGATCCcagaGATGGAGCAGTTATACTCT<br>AGACTCACTATAGAGCGCTCCGGTTTTTCTGTGCTGAACCTCAGGGGACG<br>CCGACACACGTACACGTC***ACCAGAATGGCCGAGTGGTTAAGGCGTTGG***<br>***ACTTCAGATCCAATGGATTCATATCCGCGTGGGTTCGAACCCCACTTCT***<br>***GGTA***GTCCTTTTTTTGCTTTAGTGAGGGTTAATTCAGCATGATCCAGCAG<br>GGTGGATGCCAGTCTGTGATTCTAAGGG**TATCAGCACACAATTGCCCAT**<br>**TAT** | 63% |
| 4xACE-tRNA$^{Leu}_{UGA}$ pMC insert (whole vector is 5 kb, phiC31 attB highlighted blue, phiC31 attP39 highlighted purple) | GTGCCAGGGCGTGCCCTTGGGCTCCCCGGGCGCGtgccGATGGAGCAGT<br>TATACTCTAGACTCACTATAGAGCGCTCCGGTTTTTCTGTGCTGAACCTCA<br>GGGGACGCCGACACACGTACACGTC***ACCAGAATGGCCGAGTGGTTAAG***<br>***GCGTTGGACTTCAGATCCAATGGATTCATATCCGCGTGGGTTCGAACCC***<br>***CACTTCTGGTA***GTCCTTTTTTTGCTTTAGTGAGGGTTAATTCAGCATGATC<br>CactaGATGGAGCAGTTATACTCTAGACTCACTATAGAGCGCTCCGGTTTTT<br>CTGTGCTGAACCTCAGGGGACGCCGACACACGTACACGTC***ACCAGAATG***<br>***GCCGAGTGGTTAAGGCGTTGGACTTCAGATCCAATGGATTCATATCCGC***<br>***GTGGGTTCGAACCCCACTTCTGGTA***GTCCTTTTTTTGCTTTAGTGAGGGT<br>TAATTCAGCATGATCCttacGATGGAGCAGTTATACTCTAGACTCACTATAG<br>AGCGCTCCGGTTTTTCTGTGCTGAACCTCAGGGGACGCCGACACACGTA<br>CACGTC***ACCAGAATGGCCGAGTGGTTAAGGCGTTGGACTTCAGATCCA***<br>***ATGGATTCATATCCGCGTGGGTTCGAACCCCACTTCTGGTA***GTCCTTTTT<br>TTGCTTTAGTGAGGGTTAATTCAGCATGATCCcagaGATGGAGCAGTTATA<br>CTCTAGACTCACTATAGAGCGCTCCGGTTTTTCTGTGCTGAACCTCAGGG<br>GACGCCGACACACGTACACGTC***ACCAGAATGGCCGAGTGGTTAAGGCG***<br>***TTGGACTTCAGATCCAATGGATTCATATCCGCGTGGGTTCGAACCCCAC*** | 12% |

| | | |
|---|---|---|
| | ***TTCTGGTA***GTCCTTTTTTTGCTTTAGTGAGGGTTAATTCAGCATGATCCgcaaGCCCCAACTGGGGTAACCTTTGAGTTCTCTCAGTTGGGGG | |
| 200 bp Optimized ACE-tRNA$^{Arg}_{UGA}$ CEDT (optimized ACE-tRNA$^{Arg}_{UGA}$ expression cassette bolded, TelL highlighted green, TelR yellow) | **GCGTATAATGGACTATTGTGTGCT**TTGATGGAGCAGTTA**GCCAGGGGTGTGGCCATACAGGTTTATAGTGGTTAGTAGAGACAAGTAAGTGGGTGCCCCTGTGGCGCAATGGATAGCGCATTGGACTtcaAATCCAAAGGTTGCGGGTTCGAGTCCCGCCAGGGGCGGACCTTTTCCCCATCAGTTTTTATAAACTTACACA**GCAGGGTGGATGCCAGTCTGTGATTCTCAGG**TATCAGCACACAATTGCCCATTAT** | 62% |

**Supplemental Table 1. Sequences of vectors used in this study.**
